# Origins Anywhere and Everywhere: Genome replication requires cooperative activation of thousands of origins in each cell

**DOI:** 10.64898/2026.08.17.745231

**Authors:** Linh Thuy Nguyen, Caitlyn Jean-Baptiste, Julie Gabetto, Bishal Acharya, Daniele Raimondi, Marta Radman-Livaja

## Abstract

Our model of replication dynamics in a single eukaryotic cell supplants current population-based models. Long-read sequencing reveals cooperative activation of ∼8400 closely spaced origins in every *S.cerevisiae* cell. Origins with bidirectional forks of comparable velocities are evenly distributed across the genome. The probability of early origin activation directly correlates with the average density of ORC (Origin Recognition Complex) binding motifs in a 10-20kbp region. We propose that 3D chromosome folding creates genomic pockets that act as ORC traps to increase local concentrations of pre-RC (Replication Complexes) and favor origin activation. Our results indicate that efficient replication through chromatin requires the activation of thousands of origins spanning the entire genome. This is consistent with genome instability being caused by insufficient origin activation rather than by perturbations in the replication timing program.

## Introduction

Genomes of all studied eukaryotes replicate according to a defined replication program implemented during the S-phase of the cell cycle. Cell population monitoring of genome-wide replication revealed that each genomic locus replicates at a characteristic time during S-phase ^1,2^. Replication timing of any given locus is defined as the time since the beginning of S-phase that it takes for that locus to be replicated in 50% of cells in a synchronized cell population. The Replication Program thus refers to the distribution of replication timing across the genome. Replication Programs of studied eukaryotes characteristically seem perturbed in pathological conditions associated with genome instability, like cancer^3^. A regulated replication program is therefore commonly considered as essential for the optimal functioning of dividing cells, although how and why should the replication program contribute to the preservation of genome stability is not at all understood.

All current models of replication dynamics in eukaryotic cells are derived from genome-wide mapping of replicated DNA in a population of S-phase cells and based on cell population averages ^4–7^. State of the art techniques (including NChAP developed in our group^6,7^) rely on the labeling of replicated DNA with Thymidine analogs BrdU or EdU followed by Illumina type NG sequencing. The accuracy of these models consequently hinges on correctly gauging the relationship between S-phase length at population and single-cell levels. Yeast cells are usually synchronized by arresting them in late G1 and then releasing the whole population into S-phase. S-phase duration is then defined as the period between the time of release from G1 when first cells start replicating till the time when the last cells in the population have finished replication. The duration of S-phase in the population will therefore depend on the rate of S-phase onset and the time it takes to replicate the whole genome in each cell of the population. Currently accepted models of replication dynamics assume, without experimental verification that: 1. most cells start S-phase in rapid succession, i.e. have a high cellular rate of S-phase onset, and 2. S-phase duration in single cells is equal to the duration of S-phase in the population.

Replication initiates at discrete genomic loci, called origins that are distributed throughout each chromosome. There are currently two competing models for the dynamics of origin activation, commonly referred to as origin firing. The deterministic model posits that each origin has a pre-determined firing time that depends on its genome location. Regions with early replication timing contain so-called “early” origins that fire early in S-phase, while regions with late replication timing contain “late” origins that fire late in S-phase. This implies that late replicating regions are spatially and/or physically “insulated” from early replicating regions in order to prevent replication forks from early origins to enter late replicating regions before the firing of late origins. Deterministic control of origin activation consequently assumes the existence of specialized molecular mechanisms operating in early and late regions. However, the precise nature of these molecular mechanisms is still not known ^1,8^. Furthermore, this model does not fit recent replication dynamics data that reproducibly detect the activation of both “early” and presumably “late” origins in early S-phase ^6,7^.

The deterministic model has, by now, been replaced by the stochastic model, which postulates that every known origin has some non-zero probability to be activated in any cell in the population during any given S-phase ^9–11^ . The stochastic model introduces the notion of origin activation efficiency, with all known origins firing with a locally constrained probability within a narrow time window in early S-phase. The differences in replication timing of different genomic regions are explained by different activation efficiencies (i.e. firing probabilities) of origins located in those regions. Efficient origins are believed to be activated in most cells, while inefficient origins are activated in only a small fraction of the population. Thus, regions with early replication timing are located close to “efficient” origins, and “late” regions are either replicated from inefficient origins in a small fraction of cells or by replication forks that came from more distant efficient origins.

While this model accounts for the observed low levels of early S-phase firing of origins from late replicating regions, it does not explain the “triangular” shape of the replicated DNA signal in early S-phase time points (see the NChAP lines in Figs. 1A and 7A). Assuming little cell-to-cell variability in replication fork velocity, the shape of the signal at any given locus is expected to be trapezoidal or almost rectangular if the above assumption of synchronized origin firing across the cell population, is true. The main drawback of either models, however, is that neither explains what determines the timing and efficiency of origin firing, nor what, if any, is the biological function of the replication program.

**Figure 1.**
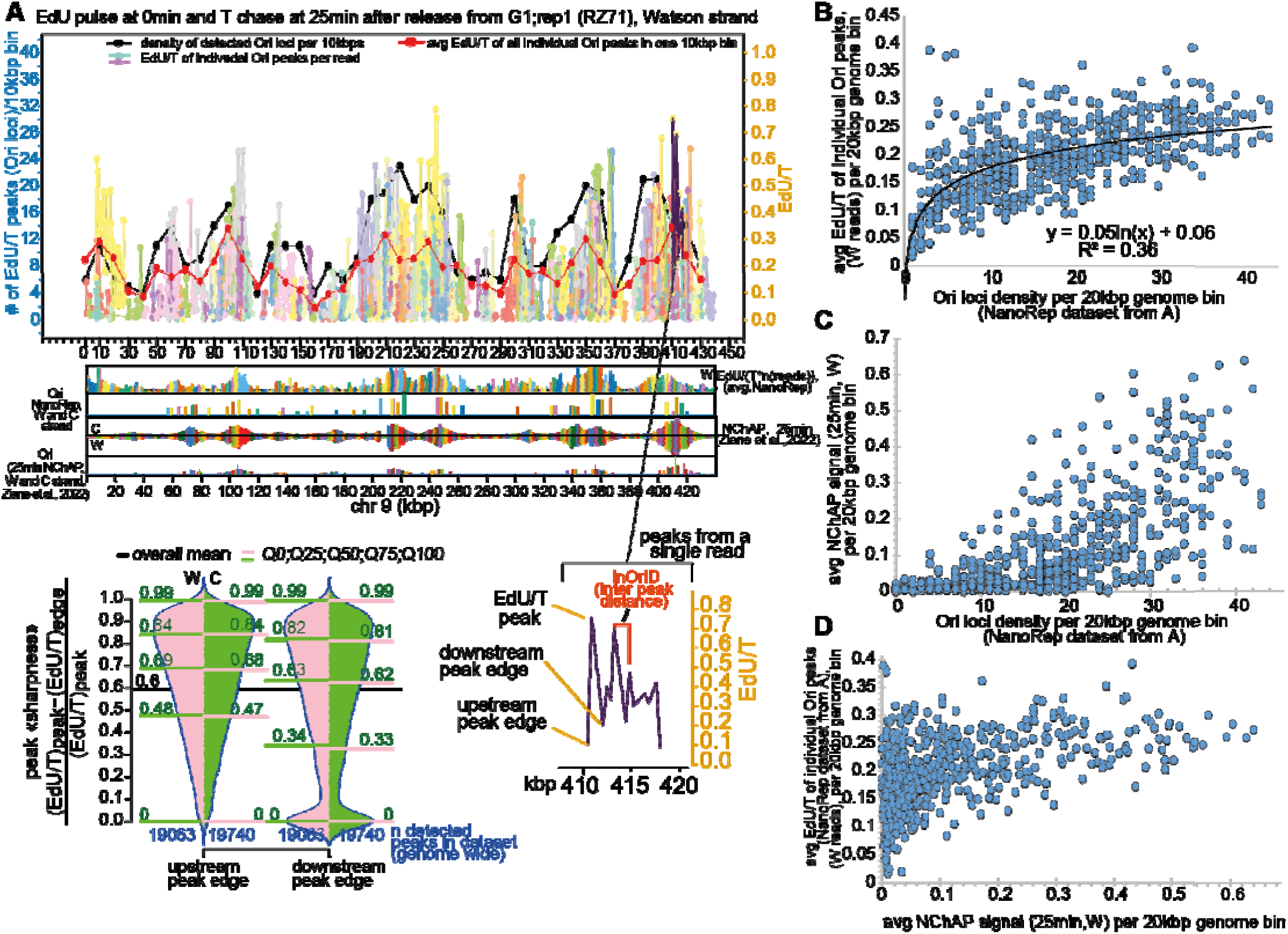
Origins from early replicating regions incorporate more EdU than origins from late replicating regions. **A. top**: <u>EdU/T of individual Ori (origin) peaks per read (pastel color palette)</u> : the distribution across chromosome 9 of EdU/T per 400bp bin (each point represents one 400bp bin, bins are contiguous and non-overlapping) in each read that contains at least one EdU. Each individual read from the EdU pulse T chase replicate 1, 25min time point (W reads only) is shown in a different color (different reads that are located far apart on the chromosome may have the same color because there are more reads than available contrasting colors and colors are assigned in order of read appearance on the chromosome). <u>Density of detected Ori loci per 10kbp</u> <u>(black line)</u>: the number of different 400bp loci (identified as EdU/T peak locations in individual reads (shown in pastel colors)) in a 10kbp genome bin (each point is one 10kbp bin, bins are contiguous and non-overlapping). Individual 400bp loci that are identified as peaks more than once in the dataset are counted only once (as one Ori locus). <u>Avg EdU/T of all individual Ori peaks in one 10kbp bin (red line):</u> average EdU/T of all Ori peaks within a 10kbp bin (each point is one 10kbp bin). Individual Oris are counted every time they appear in the dataset. Only the EdU/T of the peak-not the edges-is used for the calculation. The panels below the graph show from top to bottom: 1. the population average distribution of EdU density on chr 9 from the NanoRep dataset shown in the main graph on top (n(reads) represents all reads, including the ones not containing EdU) 2. Oris identified from 1. 3. Population average density of the NChAP signal on chr 9 from the 25min S-phase time point from^6^. 4. Oris identified from 3.**bottom:** The bean plot on the right shows the density distribution of peak “sharpness” (defined as [(EdU/T)_peak_–(EdU/T)_edge_]/ (EdU/T)_peak_=> peak “sharpness” increases as the ratio approaches 1) for all full peaks (see “EdU/T peak detection” in Materials and Methods) that were identified in the dataset. The peak and edges are defined as shown in the diagram on the right, which shows the EdU/T/400bp in one individual read from the graph in the top panel. W: Watson reads; C: Crick reads. **B-D.** Scatter plots showing the correlation between the number of identified Ori loci per 20kbp non-overlapping genome bin (defined as in A, each point is a different 20kbp genome bin) and the average EdU/T of all individual Ori peaks (defined as in A) (**B.**) or the average NChAP signal^6^ (**C.**) in that same 20kbp bin; and between the average NChAP signal and the average EdU/T of all individual Ori peaks in the same 20kb genome bin (**D**.).

The canonical *S.cerevisiae* replication origin is a ∼200bp sequence called ARS (Autonomous Replication Sequences). A functional ARS should contain an 11bp ARS Consensus Sequence (ACS, Fig. S1A) and two adjacent AT rich motifs (B1 and B) ^2^. The entire genome is believed to contain ∼300 origin centers distributed approximately every 30kbp along each chromosome^12^. The Origin Recognition Complex (ORC), which recognizes the ACS and B1 motifs, is preloaded to origins and together with the helicase Mcm2-7, cdc6, and cdt1 forms the pre-replication complex (pre-RC) during origin licensing in G1. Origin firing in early S-phase is marked by the pre-RC transition to the pre-initiation complex with the addition of DNA polymerase complexes ^10,13–16^. Thanks to the high-resolution power of (NChAP)^6,7^, we discovered that each origin center ^12^ is constituted of a cluster of closely spaced origins ^6^. We identified more than 2000 origin loci with at least one ACS motif. Only ∼15,000 of the ∼25,000 ACS motifs are found in these origins, raising the question of why yeast maintains such a large excess of apparently unused ORC-binding sites.

It goes without saying that it is impossible to infer with any confidence how genome replication proceeds in single cells, using solely population-based data. Moreover, given that as described above, several key predictions from current population-based replication models are not fully compatible with the sequencing data that these models are supposed to explain, we set out to develop a single-molecule sequencing technique for quantitative analysis of replication dynamics in single yeast cells.

## Results

### Origin firing efficiency is not determined by the DNA sequence immediately surrounding the origin peak

First, we asked whether the immediate sequence (+/-200bp) surrounding origin peaks identified by NChAP^6^ influences origin efficiency (**Fig. S1A**). Mapped origins are all located within AT rich sequences, as expected, but there are no discernable differences in the sequence composition of the ACS motif or the surrounding sequence that account for the differences in origin firing efficiency. To empirically verify this, we trained and tested five machine learning models to try to predict the firing efficiency of the 500bp origin regions centered around an ACS shown in Fig. S1A. All models failed to find any signal in the DNA sequence alone (Spearman ρ≈0, **Fig. S1B,** and **Table 1** in **Computational Methods S1** in Supplementary Materials).

Next, we tested the long held belief that chromatin organization and gene expression levels directly impact origin firing efficiencies. We looked at the localization of efficient and inefficient origins relative to the tss and RNAPol2 occupancy of the closest gene (**Fig. S1C**). More than half of all mapped origins overlap with gene promoters. This is consistent with findings showing that origins and promoters have on average a similar nucleosome architecture, consisting of a ∼200bp nucleosome depleted region flanked with well positioned nucleosomes ^17^. Even though nucleosome free promoter regions are slightly favored, a little less than half of all origins are also found within the gene coding sequence, suggesting that a promoter-like nucleosome architecture is not essential for origin activation. Furthermore, neither the localization of the origin relative to a gene promoter nor the expression level of the closest gene show any correlation to origin efficiency (**Fig. S1C**), thus invalidating the notion that chromatin organization and/or transcriptional activity are defining features of the yeast replication program.

Since population-based models fail to fully explain replication dynamics data, we developed NanoRep, a single-molecule technique based on direct detection and quantitative analysis of EdU incorporation into replicated DNA using long-read nanopore sequencing.

### 8400 origins, equally distributed between “early” and “late” replicating regions, are activated in each cell

Any model of cellular replication dynamics requires accurate estimates of both the cellular rate of S-phase onset in synchronized populations and S-phase duration in individual cells. We therefore designed two types of NanoRep time course experiments that measure S-phase progression in a synchronized cell population (**Fig. S2**). The first - an EdU pulse followed by a series of Thymidine chase steps-measures the rate at which cells start replicating after release from G1 arrest (**Fig. S2A**). The second-a series of EdU pulses after release from G1 arrest-measures the rate at which cells finish replicating (**Fig. S2B**). Since our yeast strains are haploid and genomic DNA is sequenced directly without PCR amplification, we can compute the fractions of cells that are replicating, i.e. have incorporated EdU, from the fraction of reads that contain EdU at each Thymidine chase or EdU pulse time point. The two experiments thus produce cumulative distributions of the fraction of cells that have respectively started or ended replication (**Fig. S2C**). We show that even though S-phase lasts ∼40min at the population level, individual cells finish replication in only ∼6min on average, with ∼50% of cells having replicated their entire genome within 10min (**Fig. S2D)**. This is explained by a slow rate of S-phase onset in individual cells: it takes 17min since the release from G1 arrest for 50% of cells to start replication at an average rate of 3%/min in the first 30min. So, by 25min (i.e. only a little more over the halfway mark of S-phase duration in the population) more than 50% of cells have already finished replicating. Yet, NChAP-like population-based experiments reproducibly show that replicated DNA is enriched in less than half of the genome by that point (see the NChAP profile in Fig 1A).

Hoping to resolve this apparent paradox, we mapped the distribution of incorporated EdU, expressed as EdU density (EdU/T) in contiguous non-overlapping 400bp bins, across every single read from the EdU pulse and Thymidine chase datasets. We found arrays of closely packed peaks on every EdU containing read (average read length is ∼10kbp), and mapped thousands of evenly spaced peaks across the entire genome (**Fig. 1A, Data S1**). More than half of detected peaks exhibit a more than 60% difference between EdU density at the height of the peak compared to its base (Fig 1A). The sharpness of the peaks excludes random fluctuations in EdU incorporation rates in the wake of a single replication fork as the cause for the observed high variability in EdU density along the same read. EdU/T peak profiles suggest instead that each peak marks the site of a replication initiation event, i.e. a replication origin. When cellular EdU is limiting and many origins fire nearly simultaneously, EdU is enriched near initiation sites and tapers off as forks progress, just as observed. Notably, EdU/T peak profiles from one read do not exactly align with those from another in the same region, indicating that initiation can occur almost anywhere. In regions of high peak density across reads, a handful of 400bp origin loci within a 10kb region will consistently align across molecules. These well-aligned peaks correspond to the origin clusters detected in population-averaged NChAP maps (**Fig. 1A**). The cell-to-cell “fuzziness” in origin positions means that population-based methods - even high-resolution ones like NChAP - detect only a subset of all potential origins.

The direct correlation between the density of different 400bp origin loci identified across all reads, and the EdU content of individual peaks from individual reads within a given 20kbp genome bin (**Fig 1 A-B**), reveals that individual replisomes in regions with a high density of detected origin loci tend to incorporate more EdU than replisomes from regions of low peak density. Detected peak density directly correlates with the NChAP signal (**Fig. 1C**), which means that the EdU content of individual origins in single cells is a direct measure of origin efficiency and replication timing at the population level (**Fig.1D**).

What sequence of events explains why early-replicating “origin jungles”(**Fig. 2**) show high peak density and high EdU content, while late-replicating “origin deserts” (Fig. 2) show both low peak density and low EdU? Fewer firing events in “deserts” could explain low peak density but not a lower EdU content per peak. If “deserts” truly had fewer origins, reads from these regions should show larger inter-origin distances. They do not: all regions display the same ∼1.5kb median spacing (**Fig. 2A**, top), implying ∼8400 active origins per cell (**Fig. 2A**, bottom left). Comparing this estimate with the total number of detected loci in the cell population yields an origin detection coefficient for each 20kb bin. “Jungles” are over-detected and “deserts” under-detected: ∼4000 “jungle” origins activated in one cell correspond to ∼65% of all loci detected in “jungles” at the population level, whereas only ∼30% of the ∼4000 “desert” origins that should be active in one cell are detected in the cell population (**Fig. 2A**, bottom right). Thus, “deserts” are not origin-poor at all; they contain as many active origins as “jungles”. We will nevertheless retain “jungle” and “desert” as useful qualifiers for distinguishing these two regions with different firing dynamics.

**Figure 2.**
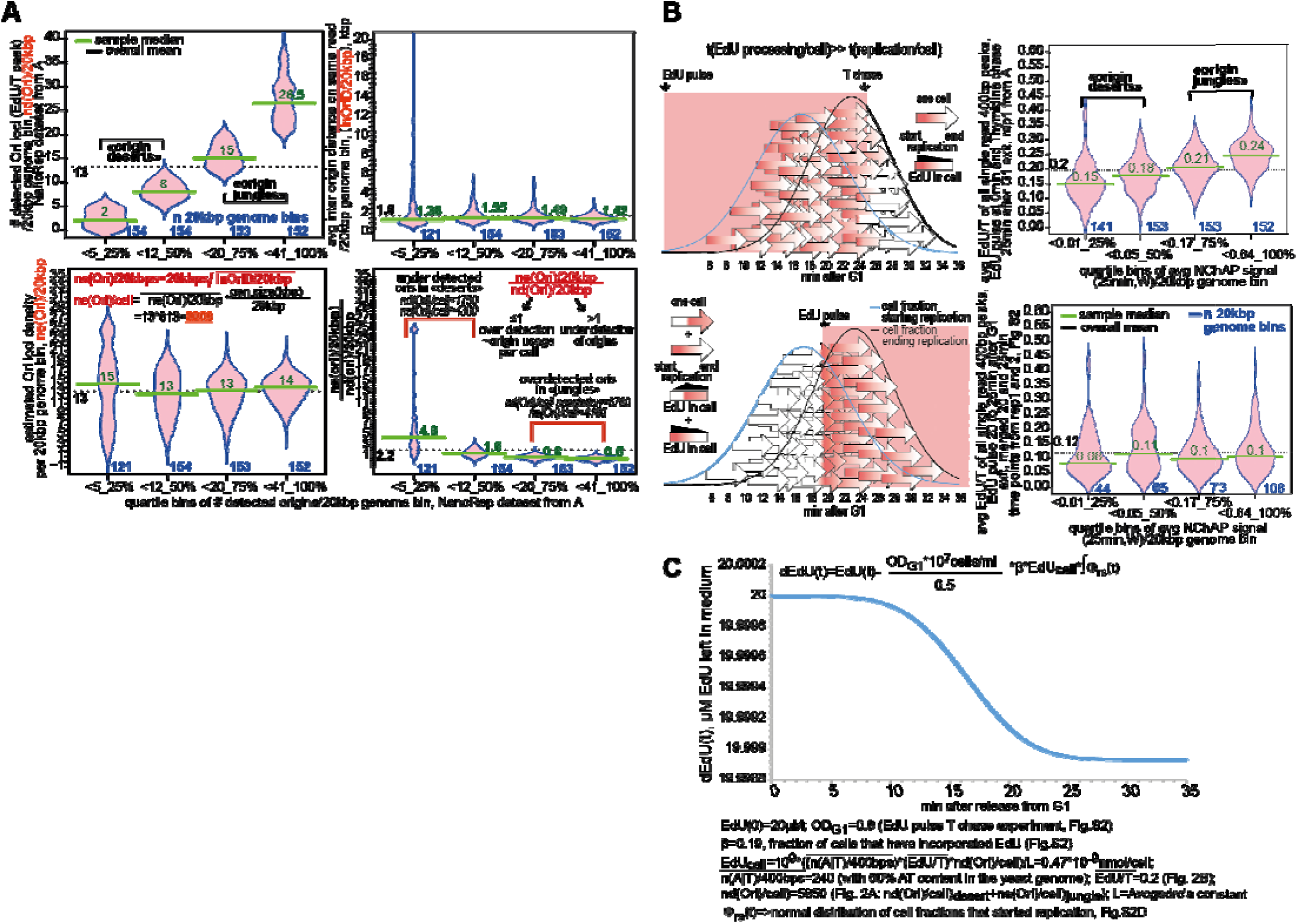
∼8400 evenly spaced origins are activated across the genome in each yeast cell. **A.** The yeast genome was divided into contiguous non-overlapping 20kbp bins. The number of Origin loci (defined as in A) detected in the dataset from Fig. 1A were counted for each bin, producing nd(Ori)/20kbp for each bin. 20kb bins were then sorted into quartiles of nd(Ori)/20kbp and the density distribution of nd(Ori)/20kbp was determined for each quartile (**top left**). We then calculated the average inter origin distance for each read in each quartile (avg(inOriD)) by measuring inOriD (defined in Fig.1A, bottom right) between each peak on each read and dividing by the number of all peaks in the 20kbp bin. Density distributions of resulting avg(inOriD) values in each quartile (**top right**) show that the average inter origin distance does not correlate with the densi y of detected origin loci. Avg(inOriD) values for each 20kb bin in each quartile were used to estimate the number of active origins per 20kbp (ne(Ori)/20kbp) bin and the number of active origins per cell (ne(Ori)/cell) (**bottom left**). Finally we used the ne(Ori)/20kbp and nd(Ori)/20kbp values in each quartile to obtain the density distributions of detection coefficients for the cell population in each quartile (**bottom right**). **B**. **left:** Diagrams illustrate when EdU gets incorporated into replicated DNA during S-phase in each cell (for a 10min duration of cellular S-phase) based on the timing of the EdU pulse and T chase after release from G1 arrest (**top**: 0min EdU pulse and 25min T chase; **bottom**: 20min EdU pulse) and the dynamics of replication start and replication end in each cell (the normal distribution curves for the cell fractions starting (blue line) and ending replication (black line) are taken from Fig.S2D). **right:** the average normalized NChAP signal avg(NCHAP) ^6^ was determined for each 20kb genome bin. 20kb bins were sorted into quartiles of avg (NChAP) and the distribution of average EdU/T per 20kbp bin (determined from values of each individual peak in the bin) was determined for each quartile of the 25min T chase time point of the EdU pulse T chase rep. 1 experiment (**top**) and the 20-25min time points of the EdU pulse rep1. and rep2. experiments, see Fig. S2 (**bottom**).**C.** Calculation of EdU usage dynamics from the media (dEdU(t)) of cell cultures during the first 35min after release from G1 arrest.

By 25 min after the EdU pulse at the time of release from G1, over half the cells have already finished replication (Fig. S2D), yet because of slow EdU processing the incorporated amount of EdU is only enough to label ∼70% of activated origins (Fig. 2A). In these conditions, early-firing origins should incorporate more EdU than late ones. Consequently, early firing origins in “jungles” have used up most of the EdU in the cell by the time origins in “deserts” start firing (**Fig. 2B**, top). We tested this by adding EdU 20min after release from G1 in the EdU pulse time course described in Fig. S2. At 20-25min after release from G1, the fraction of cells that are finishing replication (i.e. firing origins in “deserts”) should be approximately equal to the fraction of cells that are just starting replication (i.e. firing origins in “jungles”). EdU added at this point should therefore label both regions equally, as shown in **Fig. 2B** (bottom).

We have established that each yeast cell activates ∼8400 origins that are evenly spread out across the entire genome. This disproves the main postulate of the stochastic model, namely that distant forks are responsible for the duplication of late replicating regions in most cells. Origins fire in close succession, with origins in “jungles” starting on average before origins in deserts. Origin efficiency is now directly linked to the timing of origin firing in each cell and uncoupled from the frequency of activation in the cell population. In other words, the question is not anymore, “if” origins will fire in any given 20kb region - as advanced by the stochastic model-but rather “when” they will fire, which is conceptually closer to the deterministic timing model.

This new replication model emerged only because EdU is highly limiting in our system. Yeast must carry multiple copies of the human nucleoside transporter hENT1 and the viral thymidine kinase (TK) to import and process enough EdU for detection by nanopore sequencing ^18–20^. We therefore asked how much of the 20µM EdU in the medium actually enters cells. Our calculations show that <0.006% is imported and incorporated into DNA (**Fig. 2C**). Thus, even with ≥5 copies each of hENT1 and TK, EdU processing remains extremely inefficient and cannot be improved by increasing extracellular EdU.

### Cooperative activation of thousands of origins in each cell replicates 98% of the yeast genome in less than 5min

We took advantage of fast cellular EdU usage rates to build a quantitative model of replication dynamics. This analysis relies on three premises: 1. The time interval it takes for the local EdU content to decrease between point A and point B on the same DNA molecule is equal to the time interval it takes for a replication fork to cover that same distance between A and B. 2. If genome replication lasts ∼10min, the most efficient origin in the cell will fire first at the 0min time point after the start of S-phase and the most inefficient origin will fire last at the 10min time point at the end of S-phase. 3. The origins with the highest and lowest EdU content across all reads correspond, respectively, to these earliest and latest firing events.

As described in **Fig. 3A**, all we have to do now is identify the origin peak with the highest (EdU/T)max and the origin with the lowest (EdU/T)min in the dataset and calculate the average rate of EdU loss per cell (EdUl): EdUl=-[(EdU/T)max-(EdU/T)min]/10min. Across all W(atson) and C(rick) reads and time points in two EdU pulse T chase replicates, EdUl is –9.92 ± 0.03%/min, meaning forks incorporate ∼20 fewer EdU molecules per 400bp each minute. Since EdUl appears to be remarkably constant between datasets (as evidenced by the low standard deviation between datasets), we can use the average EdUl to determine the timing of origin firing of each detected origin on every single read (t_0n_) from each dataset from the EdU pulse and T chase time course experiment as explained in Fig 3A. Note that in our experimental conditions with limiting cellular EdU concentrations, a ∼10min S-phase duration/cell, and a slow S-phase onset rates (Fig. S2), the 15min, 25min and 35min T chase time points behave as biological replicates with similar cellular dynamics of EdU/T (see diagram in the top left panel of Fig. 2B).

**Figure 3.**
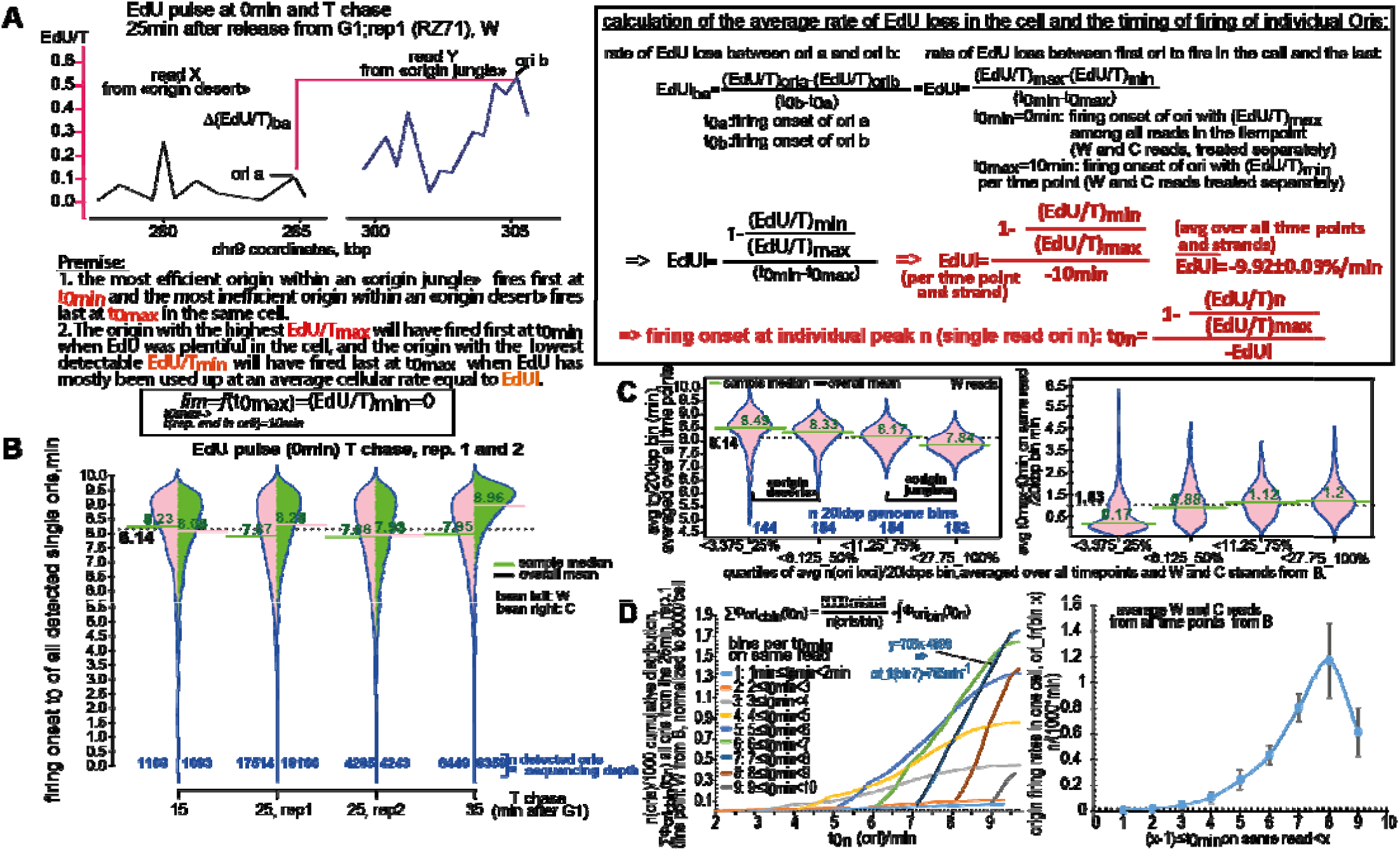
Origin firing rates in single cells reveal cooperative origin activation. **A.** Formulas used to determine the timing of firing t_0n_ of every origin peak n from all rep. 1 and 2. time points of the EDU pulse and T chase experiment from Fig. S2 are shown in the panel on the right. The variables used in the formulas are illustrated in the diagram on the left with EdU/T /400bp profiles with two reads taken from Fig. 1A. **B.** Bean plot density distribution of firing times t_0n_ for all detected origin peaks from all rep. 1 and 2 time points of the EDU pulse and T chase experiment from Fig. S2. W-Watson reads; C-Crick reads; sequencing depth see Fig. S2A. **C.** We determined the average density of Ori loci per 20kb genome bin (avg n(Ori loci)/20kbp) as in Fig. 2A but with all time points and W and C reads from rep1 and rep2 of the EdU pulse T chase experiment (from B.), and sorted genome bins into quartiles of avg n(Ori loci)/20kbp. We then determined the average firing time t_0_ for every genome bin (avg t_0_/20kbp) using t_0n_ values from B, and plotted the density distribution of avg t_0_/20kbp for each quartile of avg n(Ori loci)/20kbp (**left**). We also determined the difference in firing time between the last (t_0max_) and first (t_0min_) firing event on the same read, calculated the average difference t_0max_-t_0min_ for all reads in every 20kbp genome bin, and then plotted the density distribution of avg (t_0max_-t_0min_)/20kbp for each quartile of avg n(Ori loci)/20kbps (**right**). **D.** All Ori peaks from the 25min rep.1 time point from B (W reads) were sorted into 1min bins (from 0 to 10min) of the firing time of the first firing event (t_0min_) that happened on the same read as each Ori peak n and we determined the cumulative distribution (normalized to 8000 activated origins per cell) of firing times t0n of all Ori peaks n in each bin (**left**). The origin firing rate in one cell (n oris/min) for each t_0min_ bin determined from the slope of the linear portion of the cumulative distributions calculated as shown in the left panel gives the number of origins that fire in one cell every minute from the start till the end of S-phase (**right**). Each point represents the average firing rate from all time points, and W and C reads from B (n=7, without 35min C). Note that the origin firing rate at 9min is underestimated because of origin under detection at the end of S-phase. The error bars mark the standard deviation from the average.

Bean plot distributions of t_0n_ per time point show that origins fire on average 7 to 8min (rep3 and rep1,2 respectively) after the beginning of S-phase, with ∼90% of origins firing between 6 and 10min (**Figs. 3B and S3A**, and **Data S1**). Origins in “jungles” fire earlier on average than origins in “deserts” with a median difference of ∼20 seconds (**Fig. 3C**, left). Neighboring origins on the same DNA molecule fire in close succession, with a 0.5min or a 1min delay between the first and the last firing event in reads from “deserts” or “jungles”, respectively (**Fig. 3C**, right). The longer delay between the first and last firing event in 20kbp regions from “jungles” compared to “deserts” is consistent with the t_0n_ density distributions from **Fig. 3B**, and suggests that firing kinetics of nearby origins are cooperative, meaning that origins in “deserts” that fire later in S-phase should fire at a higher rate than origins in “jungles” that fire earlier. Plotting the cumulative distribution of t_0n_ per bin, grouped by the earliest firing event on the same read (t_0min_), and extracting firing rates from the slope of each curve (**Fig. 3D**, left), shows a clear pattern: origin-firing rates on a given DNA molecule rise steadily as the first firing event in that region approaches the 10min mark and the end of S-phase (**Fig. 3D**, right). This means that activation events at neighboring origins are not independent of each other, i.e. past origin firing events stimulate the activation of nearby origins that have not yet fired.

Next, we used the average cellular rate of EdU depletion (EdUl) to calculate DNA synthesis rates at individual replication forks as described in **Fig. 4A**. We find an average fork speed of 508 bp/min for EdU pulse T chase rep1, 2 time points (Fig. 4B, Data S1), and 365 bp/min for rep. 3 time points (**Fig. S3B**). These values are comparable to fork speeds determined from NChAP data ^6^, thus confirming our previous observation based on population averages that most replication forks progress 2 to 4 times slower than previously thought^9^.

**Figure 4.**
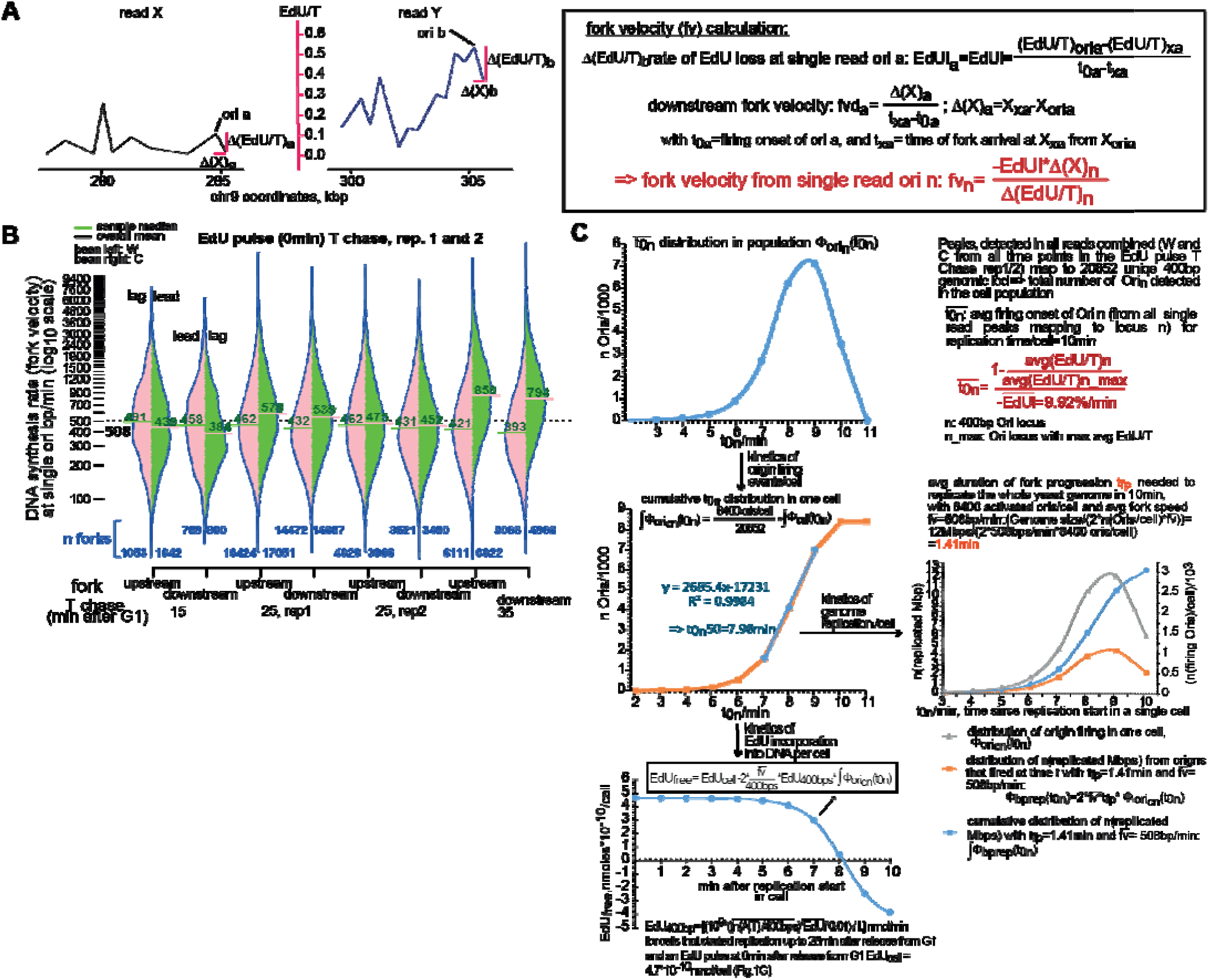
Dynamics of genome replication in single cells. **A.** Formulas used to determine the velocity of individual forks fv_n_ from every Ori peak n detected in all rep. 1 and 2 time points of the EDU pulse and T chase experiment from Fig. S2 are shown in the panel on the right. The variables used in the formulas are illustrated in the diagram on the left with EdU/T /400bp profiles with two reads taken from Fig. 1A. **B.** Bean plot density distribution of fork velocities fv_n_ from all detected origin peaks from all rep. 1 and 2 time points of the EDU pulse and T chase experiment from Fig. S2. W-Watson reads; C-Crick reads; lag-lagging strand replication; lead-leading strand replication. **C. top:** The distribution of the average firing time (avg(t_0n_)) for every 400 bp Ori locus mapped from all datasets in Fig. 3B is calculated as described on the right of the graph. The graph in the **middle panel on the left** shows the cumulative distribution of the avg(t_0n_) distribution from the top panel, normalized to the estimated number of active origins per cell (Fig. 2A). This distribution function is used to simulate the dynamics of genome replication (**middle right**) and the dynamics of EdU usage (**bottom left**) in a single cell.

By combining the average fork velocity with the timing of firing onset, we can infer genome-replication dynamics in single cells (**Fig. 4C**). Origin peaks from all datasets (**Fig. 3B**; Data S1) map to 20,852 unique 400bp loci. We found at least one origin in every 40kbp region of the genome, except at the end of chromosome 13 (**Data S2**). Using the average firing onset (avg(t_0n_)) per locus, we find that 98% of origins fire between 5 and 10min after S-phase entry, with 60% firing between 7 and 9min (**Fig. 4C**, top). A cumulative distribution normalized to the estimated 8,400 active origins per cell (**Fig. 2A**) shows that half fire by 8min, with ∼2,700 firing events/min from 7 to 9min after the beginning of S-phase (**Fig. 4C**, middle left). Rapid activation of thousands of closely spaced origins enables fast genome duplication despite slow forks. Each fork needs to run for only ∼1.41min and traverse ∼700bp to complete replication in 10min. The simulation of genome replication dynamics in **Fig. 4C** (middle right) shows that 98% of the genome is replicated in the second half of S-phase. The first half of the genome requires ∼8min, while the remaining half is completed in ∼2min thanks to cooperative origin activation.

### Regional ACS density influences the timing of origin firing

In each cell, the genome half containing origin “jungles” is replicated from ∼4000 origins firing during the first four quintiles of S-phase, while the “desert” half is replicated from ∼4000 origins activated mostly in the final quintile. Thus, replication in jungles and deserts is qualitatively the same: both halves are duplicated in ∼1.5-kb segments from an equal number of active origins. The under-detection of late origins is consequently an experimental artifact caused by limiting cellular EdU and slow rates of S-phase entry in a synchronized cell population. In reality, the number of active origins in early and late replicating regions is the same, which is probably why no one had been able to explain in a way that made biological sense why population based data made it look like some genomic regions had more active origins than other parts of the genome. What remains is understanding why “jungle” origins consistently fire earlier than “desert” origins, and whether this conserved order of origin activation really does contribute to genome stability.

The ORC-binding ACS motif has long been viewed as essential for origin function. There are ∼25,000 ACSes genome-wide, about one every 800bp in 98% of the genome. Still, only 45% of the 20,852 identified 400bp origin loci contain an ACS (**Data S2; Fig. 5A**), suggesting that an ACS in a 400bp window is not required for initiation in that same window. Likewise, constitutive nucleosome depletion typically found at promoters and ends of genes ^21^, another feature thought to favor origin activation ^17^, is not necessary for replication initiation, as only 38% of origins are located immediately upstream of a transcription start site (tss) or immediately downstream of a transcription termination site (tes). The discrepancy between the low overlap in constitutive nucleosome depletion and the purported requirement for low nucleosome density at activated origins can however be readily explained by ORC’s nucleosome eviction capabilities ^22,23^. ACS presence also does not promote early firing: although 81% of loci fire first in a given 20kbp region in at least one cell, only 44% of those contain an ACS (**Fig. 5A**, top). ACS number or the presence of a nucleosome depleted region (NDR) also fail to distinguish “jungles” from “deserts”, which show similar distributions of either feature (**Fig. 5A**, bottom panels 1 to 4 from the left). Overall, these results reinforce our earlier conclusion that local DNA sequence near initiation sites does not determine origin efficiency at the population level (Fig. S1).

**Figure 5.**
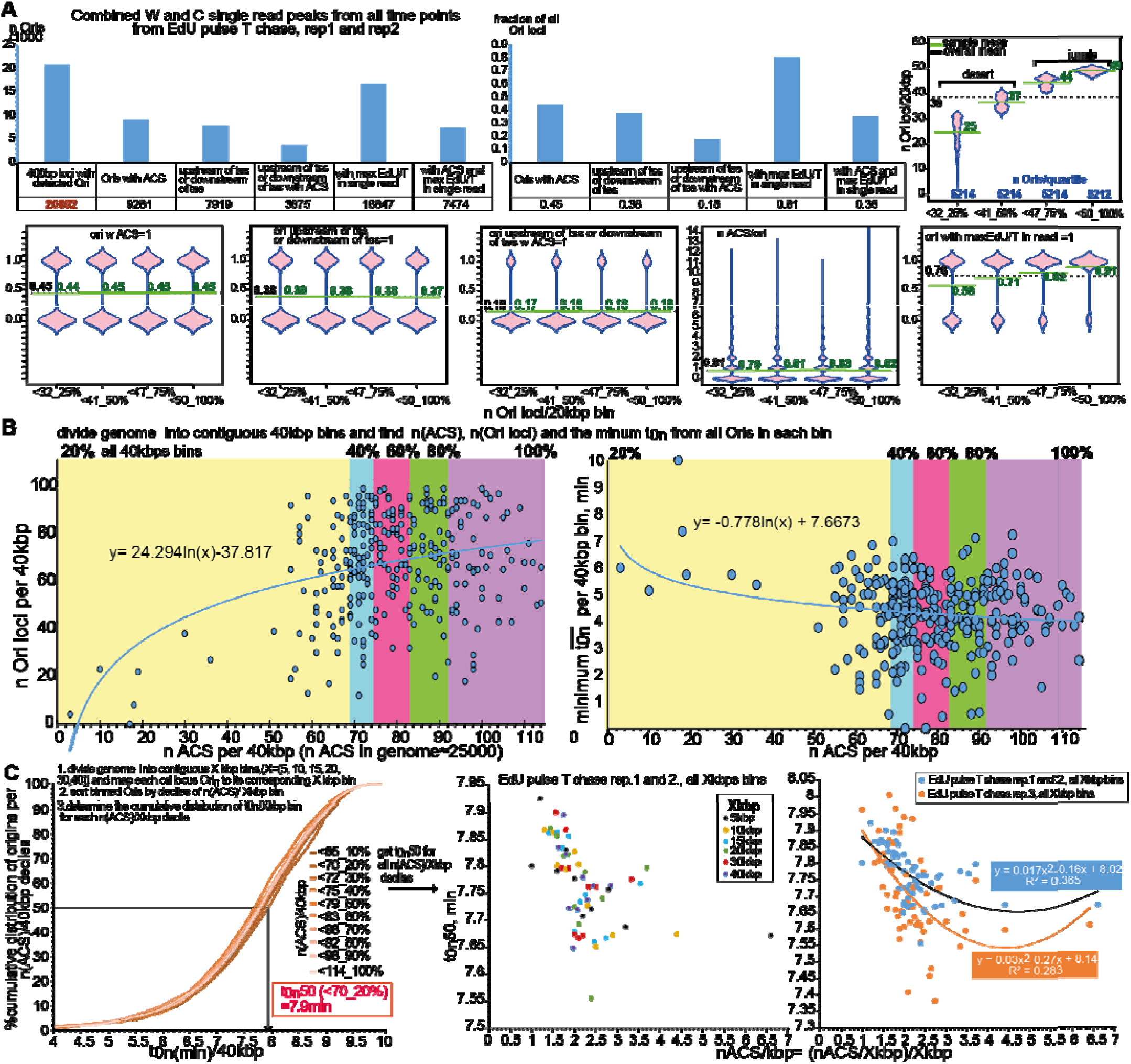
Average regional ACS density directly stimulates origin firing in early S-phase. **A.** The bar graphs show the proportion of all detected 400bp origin loci (defined as in Fig. 1A) that also have the indicated feature. The bean plots show the distribution of ori loci with the indicated features across 20kbp genome bins sorted into quartiles of density of detected ori loci. “Ori with max EdU/T in single read” means that a given Ori locus contained a peak with the highest EdU/T out of all the peaks on the same read in at least one read in the entire dataset. max(EdU/T) on the read means that that locus fired first in the cell from which the read was extracted in the genome region covered by that read. tss and tes annotations are from ^42^ **B.** The genome was divided into contiguous non-overlapping 40kbp bins. We counted the number of ACS motifs (i.e. the number of occurrences of every possible ACS consensus sequence (see Fig. S1A)) in each bin and plotted it against the number of ori loci per 40kb bin (**left**) or the smallest average firing time t_0n_ (calculated as in Fig. 3C) i.e. the minimum t_0n_ out of all the Ori loci in each bin (**right**). Each point in both graphs represents one 40kbp genome bin. The colored blocks on the graph represent the quintiles of n(ACS)/40kbp across the genome. **C.** The genome was divided into contiguous 5, 10, 15, 20, 30 or 40kbp bins and the bins were sorted by deciles of ACS density (n(ACS)/Xbp). We then determined the cumulative distribution of average firing times t_0n_ of all 400bp ori loci n (calculated as in Fig. 4C) in each X kbp bin within each nACS/Xbp decile (left). The mean firing times t_0n_50 for each n(ACS)/Xkbp decile were then determined from the cumulative distributions and plotted against n(ACS)/kbp (middle and right panels) to show the inverse correlation between origin firing times and local ACS density, i.e. origin tend to fire earlier in regions with higher ACS density.

The only consistent difference between “jungle” and “desert” origins is the chance that a locus fires first within its 20kb region. In early-replicating “jungles”, any locus has an 87% probability of firing first in at least one cell, whereas in late-replicating deserts this drops to 65% (**Fig. 5A**, bottom panel, 5^th^ from the left). This reflects the more cooperative, later firing in “deserts” (**Fig. 3D**). In “jungles”, any locus can initiate ∼1 min before others (Fig. 3C). In deserts, strong cooperativity causes origins to fire almost synchronously, reducing the likelihood that any single origin fires first.

What then, increases the probability of very early firing in origin jungles? Although an ACS need not lie within 400bp of an initiation site (**Fig. 5A**), we conjectured that regional ACS density should still influence ORC binding. We therefore tested whether ACS levels averaged over larger windows (from now on referred to as ACS density) are more predictive than ACS presence at the initiation site. Dividing the genome into contiguous 40kb bins and counting ACSes revealed a clear correlation between ACS density and the abundance of origin loci (**Fig. 5B**, left): bins with 70-90 ACSes typically contain >60 origins, whereas bins with <40 ACSes never exceed 40. Because the local density of identified origin loci correlates with firing time (**Fig. 3C**), this pattern extends to the minimum timing of firing at individual origin loci. Regions with <40 ACSes/40kb never fire in the first 5min of S-phase, while bins with 70-90 ACSes/40kb are the only ones to contain origins firing in the first minute (**Fig. 5B**, right).

Our analysis in **Fig. 5C** shows that there is an inverse correlation between the average ACS density in 10 to 20kbp regions and the mean origin firing time in the same region. Regions with more than two ACSes/kbp reproducibly fire ∼10 seconds before regions with an average ACS density between 1 and 2 /kbp (**Fig. 5C**), consistent with our initial hypothesis that the local average ACS density influences the timing of origin firing, which can nevertheless happen anywhere within that broader region irrespective of specific ACS locations.

Consistent with cooperative origin firing kinetics (**Fig. 3D**), the probability that an origin locus (out of all ∼21000 mapped loci) will fire first in a given region increases as S-phase progresses and the local density of active origins in adjacent regions increases (**Fig. 6A**, top left). Furthermore, the exponential increase in the genome-wide cellular rate of first firing events per 10kb (avg. read length) in the first half of S-phase, is consistent with cooperative activation of origins located more than 10kb from each other (**Fig. 6A**, right). As expected, origins that fire first and earlier in S-phase come from regions with a higher average ACS density (n_50_(ACS)/kbp, **Fig. 6A**, bottom left). Subsequent firing events within each 10kb region closely follow the first one, and the pace of origin firings accelerates as S-phase progresses. Four additional origins fire within 4min, 2.5min or 1.5min of the first if the latter respectively fires between 0 and 6min, 6 and 8min, or 8 and 10min after the beginning of S-phase (**Fig. 6B**). The firing probability rate increases exponentially as S-phase progresses and more origins are activated in the surrounding regions (**Fig. 6C**). Thus in a given genomic region, the timing of origin firing is inversely correlated to firing rate, and origins in regions with higher ACS density have a higher probability of firing early in S-phase.

**Figure 6.**
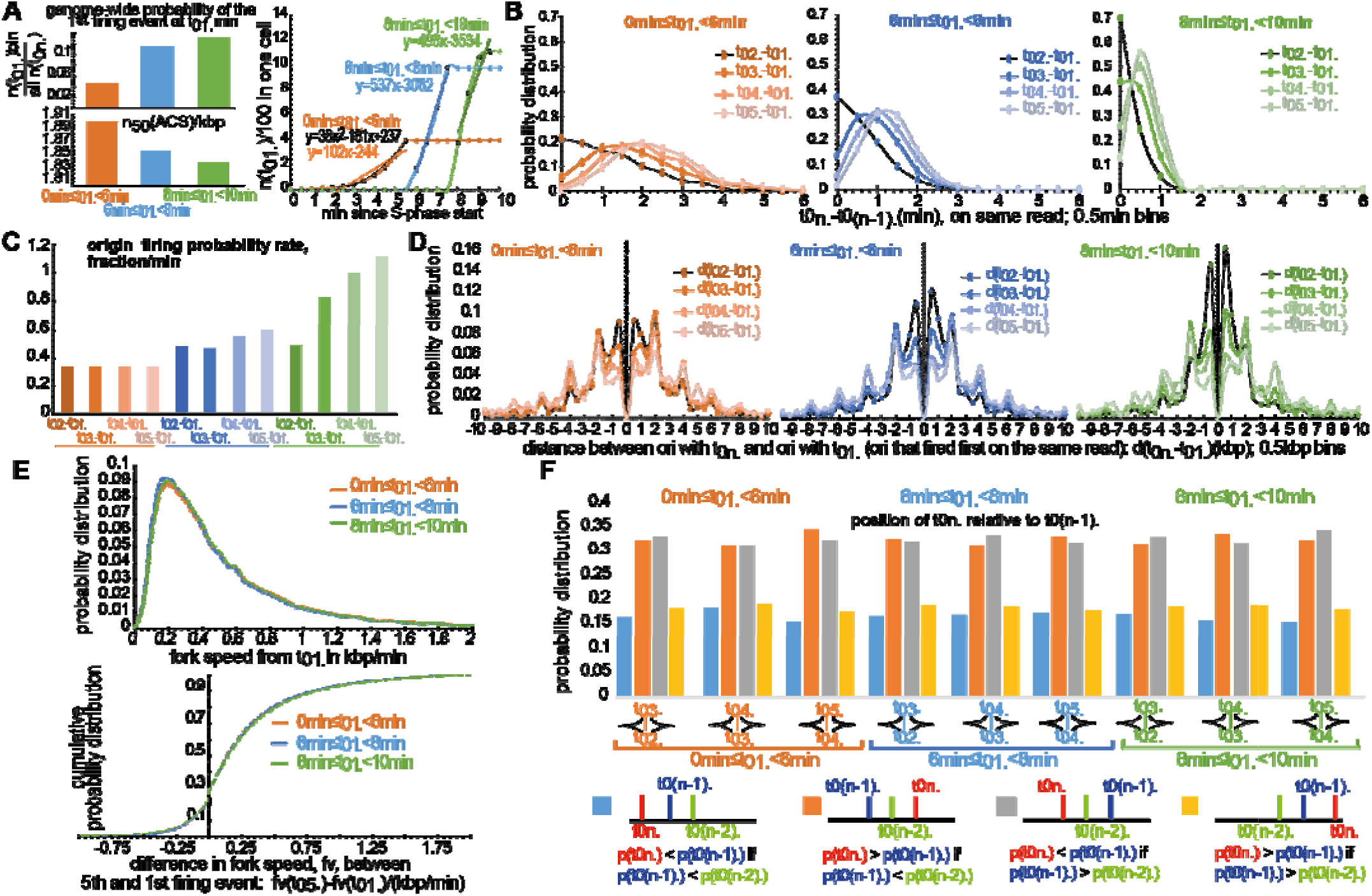
Replication proceeds through a cooperative cascade of closely packed initiation events. **A. top left**: We found the ori peak with the smallest (earliest) t_0n_ (calculated as in Fig. 3A) on every read from all datasets from rep. 1 and 2 of the EdU pulse T chase experiment. This is the ori that fired first (t_01._) on a given read. We then counted the number of first firing events that happened in the first half (0 to 6min), the 3^rd^ quintile (6 to 8min), or the last quintile (8 to 10min) of S-phase (n(t_01_)bin) and divided by the total number of firing events that happened any time in S-phase across the cell population (all n(t0n)=47199 (mapping to 20852 unique 400bp loci), from all datasets from rep. 1 and 2 of the EdU pulse T chase experiment).This gives the probability for any detected ori locus to fire first in a given ∼10kb genomic region (average read length is 6±5kb) at specific times during S-phase. **bottom left**: We determined the cumulative distribution of ACS densities (n(ACS)/kbp, calculated as in Fig. 5C.) of all reads within each t_01_ bin from the top panel, and calculated the mean ACS density (n_50_(ACS)/kbp) from the cumulative distributions. **right**: cumulative distributions of first firing events t_01._ (calculated as in the top left panel) normalized to the total estimated number of active origins per cell (8400) during S-phase: first half-orange, 3^rd^ quarter-blue, and last quarter-green. The linear and polynomial fits show the firing rates of earliest firing events per >=10kb region during S-phase. **B.** The graphs show the distribution of time intervals between the first firing event (t_01_.) and subsequent firing events on the same read (t_0n._, n=2, 3, 4 or 5) at different times during S-phase (same bins and dataset as in A.). **left**: first half, **middle**: 3^rd^ quintile, **right**: last quintile**. C.** The probability of firing events after the first firing event on the same read per min is calculated from the slopes of the linear part of the cumulative distributions derived from distributions from B. **D.** The graphs show the distribution of distances between the position of first firing event (t_01_., centered at 0 on the graph) and the positions of subsequent firing events on the same read (t_0n._, n=2, 3, 4 or 5) at different times during S-phase (same bins and dataset as in A. and B.). **left**: first half, **middle**: 3^rd^ quintile, **right**: last quintile**. E. top**: Distribution of fork velocities (averaged between upstream and downstream forks) from all first firing events from S-phase bins from B. **bottom**: Cumulative distribution of differences in fork speeds (averaged between upstream and downstream forks) between the forks issues from the fifth firing event and forks issued from the first firing event on the same read from the same S-phase bins as in B. **F.** Probability of positioning p(t_0n._) of a firing event t_0n._ (n=3, 4 or 5) relative (upstream or downstream) to the two preceding firing events t_0(n-1)._ and t_0(n-2)._

How far does the stimulatory effect of the first firing event in a 10kbp region extend? **Fig. 6D** shows that the 2^nd^–5^th^ firing events on the same DNA molecule remain tightly clustered, rarely occurring more than ±5kb from the first. Early and late S-phase regions show similar patterns, but late regions display tighter clustering: the 2^nd^ and 3^rd^ events occur within ∼500bp of the first, compared to ∼2kb in early S-phase. Because fork velocities originating from the first firing event remain constant throughout S-phase (**Fig. 6E**, top), this reflects a faster firing rate later in S-phase - forks have less time to advance before nearby origins activate. High local fork density also accelerates forks from later firing origins on the same DNA molecule. About 70% of forks from the 5^th^ firing event in a 10kb region are faster than those from the first, and half are at least 150 bp/min faster (**Fig. 6E**, bottom).

New firing events occur equally upstream and downstream of the first event, as shown by the symmetric distributions in **Fig. 6D**. When firing events on the same read are centered around the first, each subsequent event is about twice as likely to appear on the opposite side of the previous one (**Fig. 6F**). Thus, origins tend to fire in an alternating pattern at regular intervals on either side of the first-firing origin, progressively moving outward until they encounter forks arriving from neighboring 10-20kb regions.

### The “topological protein trap” model of replication dynamics in single yeast cells

Our analysis shows that replication can initiate anywhere in the genome. Although 98% of origins fire in the second half of S phase, regions with higher ACS density (≥2/kb) fire slightly earlier (∼10 s) than those with lower density (1–2/kb). Within any 10–20 kb region, the first firing event positions subsequent ones, which occur at regular 1–2 kb intervals on either side of the first one. Firing times of individual origin peaks allow us to compute firing probabilities for every 400 bp locus (**Fig. 7A, Data S3**). Because these probabilities rely on detected origin peaks - 20,852 out of the 30,175 possible non-overlapping 400bp loci (**Fig. 5A**) - we underestimate firing late in S-phase due to EdU depletion (**Figs. 2A**, **4C**), whereas probabilities for loci active in the first 75% of S phase are reliable.

In early replicating “jungles,” firing probability crests in the third quarter of S phase. Since replication proceeds through dense cascades of origin firing events rather than elongation from a few widely spaced origins, every 400 bp locus should fire in at least one cell by S-phase completion, giving each locus a cumulative firing probability of 1 in the cell population. Loci with low probability of firing in early S-phase must therefore fire in the final quarter, allowing inference of firing probabilities in late replicating “deserts” (**Fig. 7A**). Notably, even loci with a higher probability of early firing actually fire late in most cells. They simply fire early in a subset of them. Conversely, “late” origins fire almost exclusively at the end of S phase. ACS density therefore determines which regions have a higher chance of early firing, but cooperative activation is the main driver of origin firing later in S phase. Population-level replication timing is therefore shaped largely by widespread and exclusively late firing in late-replicating regions.

**Figure 7.**
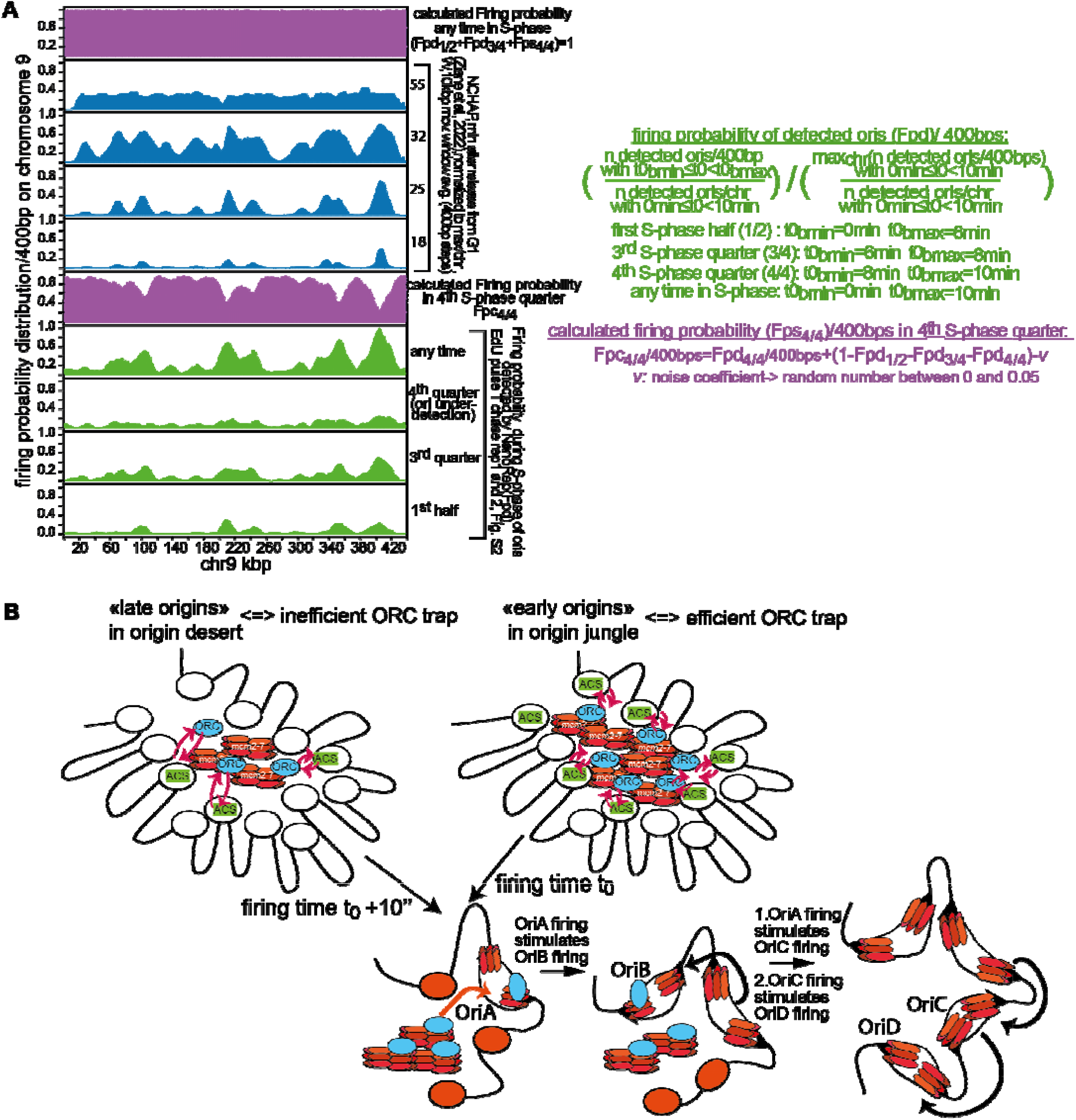
The “topological protein trap” model of origin firing dynamics. **A.** The probability Fpd that replication will initiate at a given 400bp genomic locus at a specific time in S-phase in any cell in the population (shown in green) is determined by counting all origin peaks from all datasets from rep. 1 and rep. 2 of the EdU pulse and T chase experiment (see Fig. 3B) that map to a given 400bp locus and have a firing time t_0n_ within the first half, or 3^rd^ or last quarter of S-phase. The obtained count number for each 400bp locus in each S-phase bin is then divided by the total number of detected origin peaks per chromosome that fired any time in S-phase (n(t0n)=47199 for all chromosomes together, as in Fig. 6A). Each one of these ratios for each 400bp locus is then normalized to the maximum ratio on the same chromosome as shown next to the figure. The sum of Fpd profiles in the first half, and 3^rd^ and last quarters of S-phase gives the probability that a given locus will fire any time in S-phase. The calculated firing probability profile for the last S-phase quarter takes into account the under-detection of origin firing at the end of S-phase and starts from the assumption that the Fpd for each 400bp locus firing in any cell, any time in S-phase should be 1 (shown in purple). NChAP profiles are shown in blue. **B.** Illustration of the topological genome trap model. Genome “pockets” with high ACS density are efficient ORC traps with high concentrations of ORC and Mcm2-7. Origins in efficient ORC traps consequently tend to fire earlier in S-phase at time t0. Conversely, origins in inefficient ORC traps with low ACS density fire later at time t0+10” on average. Once an origin Ori A, which could be anywhere on the DNA within the pocket, has fired it stimulates the firing of origins in the immediate vicinity and starts a cascade of origin firing within the pocket. Cdt1 and cdc6 are not included in the illustration for the sake of clarity.

The measured probability of firing any time in S phase is <1 for most loci (**Fig. 7A**) because NanoRep detects only a small fraction of late firing events, due to the fact that in synchronized cells during the first ∼30 min after G1 release, more cells enter S phase than finish it, and EdU is limiting (**Fig. 2B**). Thus, the measured cumulative firing probability in single cells matches the NChAP signal at 32 min, and the NanoRep firing-time distributions for the first half and third quarter of S phase resemble NChAP profiles at 18 min and 25 min, when most cells are still in early S phase (**Fig. 7A**).

NChAP enriches for any locus from any cell that has incorporated sufficient amounts of EdU between G1 release and cell collection, so late-replicating regions become labeled as much as early regions only after ∼45 min, once remaining S-phase cells have imported enough EdU. Because individual forks travel only ∼700 bp and thousands of origins fire across the population, origin firing profiles from NanoRep analysis (**Fig. 7A**) approximate well population level replication dynamics captured by NChAP, even though NChAP also includes elongation.

The fact that we are able to reconstruct NChAP profiles derived from populations in early S-phase, by solely using origin firing times from single cells validates our model and, unlike current models of replication programs, fully explains population-level replication timing data.

Next, we examined more closely the relationship between local ACS density and origin firing dynamics across chromosomes. Consistent with **Fig. 5B**, genome-wide average firing probabilities during the first three quarters of S-phase increase proportionally with ACS density (**Fig. S4A**). This relationship is even stronger within chromosome bodies, where ACS density has roughly twice the effect observed genome-wide (**Fig. S4B**). In contrast, chromosome ends show the opposite pattern: higher ACS density correlates with lower firing probability in early S-phase (**Fig. S4C**), suggesting an additional mechanism that actively delays subtelomeric and telomeric origin firing to late S-phase. Such a negative “telomere effect” on early origin firing suggests that replicating chromosome ends after the rest of the genome, increases the fitness of yeast cells.

The Sir complex is the most likely factor delaying origin firing at chromosome ends. Known for repressing silent mating-type loci on chromosome 3 ^24^, it also binds strongly to subtelomeric regions ^25^, where its role remains unclear. Sir recruitment depends on a variety of combinations of Orc1p, Rap1p, and Abf1p motifs. The specific arrangement of these motifs found in most subtelomeric regions and silent mating-type loci is particularly favorable to stable Sir assembly. Consequently, the occupancy of Sir3 (a subunit of Sir) is at least six fold higher at chromosome ends than in chromosome bodies with similar ACS densities (Fig. S5A, Data S3). Such high Sir3 levels correlate with reduced probability of early firing at chromosome ends (Fig. S5B–C), whereas lower Sir3 occupancy in chromosome bodies has a weak stimulatory effect on early firing (Fig. S5C). These results support a model in which strong Sir enrichment postpones replication of chromosome ends until late S phase, probably by competeing with ORC for chromatin binding.

Although ACS density promotes, on average, early origin firing genome-wide (Fig. S4), it does not fully account for locus-specific variability in early activation probabilties across chromosomes (**Fig. 7A**). We therefore used the average chromosomal ACS density/10kbp as a baseline firing-probability landscape and incrementally added biologically relevant chromatin features to better match early S-phase firing profiles from NanoRep and NChAP (**Fig. 7A**). Our aim was to identify which features shape the distribution of early- and late-firing regions along each chromosome. We first multiplicatively corrected the average ACS density per 10 kb using ORC ChIP-seq peaks ^26^, Hi-C contact matrices ^27^, Sir3 occupancy (Fig. S5, Data S3, ^28^), and RNAPol2 occupancy ^6^ (**Figs. S6 and S7** and **Computational Methods S2** in Supplementary Materials). We then trained three linear and nonlinear machine-learning models to predict early S-phase firing probabilities (**Computational Methods S3** and **Figs. S8 to S11** in Supplementary Materials), comparing outputs to firing probabilities from the 0–6 min (1^st^ half of S-phase) and 6–8 min (3^rd^ quarter of S-phase) NanoRep profiles (Fig. 7A; Data S3; Figs. S9–S11). Both modeling methods show that ORC binding proximity to a genomic region in 3D space is the strongest predictor of early origin firing in that region. This convergence motivated the ORC “trap” model of genome-wide origin-firing dynamics described below.

Chromosomes fold into 3D TADs whose distribution correlates with replication timing ^31,32^. Consistent with our computational modeling results (Figs. S6–S11), we propose that 3D folding generates 10–20 kb pockets with variable ACS densities that act as ORC traps in G1 and become Replication Domains in S phase **(Fig. 7B**). ORC diffuses freely between pockets but lingers in those with higher ACS density, increasing local ORC concentration and recruiting more Mcm2-7, Cdt1, and Cdc6. Continuous ORC turnover on ACS binding sites thus maintains a stable pre-RC pool in each pocket.

The topological protein trap model thus explains genome-wide replication timing differences (**Fig. 7**). Pockets with a higher ACS density trap more pre-RCs, increasing ON rates and promoting early firing, whereas pockets with lower ACS density retain fewer pre-RCs and fire later, as seen in origin “jungles” or “deserts”, respectively (**Fig. 3C**). Since pre-RCs share each pocket with many other proteins, ON rates also depend on local concentrations of competing factors that influence pre-RC diffusion in and out of the pocket, as well as pre-RC’s access to chromatin, as seen for the Sir complex at chromosome ends (Fig. S5).

Our modeling approaches not only underscore the role of 3D chromosome conformation but also confirm the inhibitory effect of telomere proximity on early origin firing (Figs. S4–S5) and uncover a positive effect of centromere proximity (Figs. S6–S7, S9–S10). The centromere effect likely stems from the tight clustering of yeast centromeres near the spindle pole body ^29,30^, which – according to our ORC trap model-should create a shared pool of pre-RC complexes and stimulate early firing in pericentromeric regions. Together, these telomere- and centromere-associated effects support the idea that selective pressure to replicate chromosome bodies before chromosome ends drove yeast to evolve mechanisms that delay firing near telomeres while advancing it within chromosome interiors.

## Discussion

We now have a fundamentally different picture of single-cell replication than the one derived from population-based data. Each yeast cell activates ∼8,400 origins in rapid succession, spaced at regular 1-2kb intervals across the genome. There are no inefficient origins that are activated only in a minority of cells. Dense, rapid origin activation enables complete genome replication in just 5-10min, about 3-4 times faster than previously inferred from population-averaged measurements.

As in deterministic timing models, regions in the chromosome body, previously labeled “early” regions, still fire slightly before “late” regions, but in single cells these differences are on average under 1 min, raising doubts about any biological significance of the replication timing program for ∼90% of the genome located beyond chromosome ends and pericentromeric regions. As shown in Figs. S2, 1 and 2, the apparently long delay between early and late replication seen in population data stems from misinterpreted experimental conditions: limiting EdU, the much shorter S-phase in individual cells compared to S-phase length in the population, and the slow staggered release from G1 arrest in single cells. In reality, with the exception of chromosome ends, there are no meaningful qualitative differences between “late, desert” regions and “early, jungle” regions. Each chromosome is partitioned into contiguous ∼15kb Replication Domains (RDs), each replicated locally by equivalent numbers of origins whose forks move relatively slowly (300–500 bp/min) and individually cover only ∼700bp on average.

Our analysis suggests that for 90% of the genome (the part located within chromosome bodies), the replication timing program is at its core largely a side effect of the variability in ACS density along yeast chromosomes. This variability likely reflects evolutionary pressures that balance efficient replication with constraints imposed by the transcriptional program. In this framework, the replication program is incidental: local ACS density only needs to be high enough to permit multiple origins to fire within a relatively small genomic region and a short time window. Thus, the on average small timing differences between regions do not reflect a functional replication-timing program but are instead evolutionary byproducts of how ACS density patterns were shaped by transcriptional requirements.

Except for concerted late and early replication at chromosome ends and pericentromeres, which is probably beneficial, the replication program for the remaining 90% of the genome appears to have no obvious biological function. Still, the reproducible order of origin firing is indicative of underlying mechanisms that probably govern all genomic processes in eukaryotes.

In our “topological protein trap” model (**Fig. 7B**), efficient licensing does not require long-lived ORC–DNA binding: the ON rate simply must be faster than ORC diffusion out of the pocket. Concomitantly, a moderately fast ORC OFF rate facilitates DNA access to transcription factors that are also present in the pocket. The constant exchange of pre-RCs and transcription factors on DNA thus supports both origin licensing and transcription. More broadly, the model reflects the crowded nuclear environment, where the efficacy of genomic processes hinges on conditions that increase the probability of productive encounters between proteins and their DNA targets ^33^. The uneven distribution of DNA-binding motifs, organized into 3D pockets, acts as a local “crowd controller” that retains some proteins and allows others to pass, ensuring optimal conditions for replication and transcription.

The trap model uncouples ORC binding to ACS sites from pre-RC activation and origin firing. Like other DNA-binding proteins, ORC can bind anywhere on DNA but lingers longer at consensus motifs. Thus, origins do not need to fire only within 400bp of an ACS (**Fig. 5A**). Firing can occur anywhere within a pocket where pre-RCs accumulate, because activation requires only transient binding near an AT-rich site. Mammalian ORCs, which lack strict sequence specificity, preferentially bind sequences prone to forming G-quadruplexes ^34^, which are frequently located at known mammalian origins ^35^. Thus, the trap model is probably also applicable to mammals, with G4-rich regions (instead of ACS-rich ones) acting as ORC retention pockets.

Our model also accounts for the strong correlation between replication timing and higher-order chromosome folding observed across eukaryotes ^31,32^. Regions that fold into pockets with higher ACS density accumulate more pre-RCs and fire earlier, while pockets with lower ACS density fire later. Thus, replication timing emerges from the combined effects of 3D folding and 1D motif distribution rather than from an active regulatory program.

The protein-trap model integrates two core architectural features: variation in the linear density of DNA-binding motifs and 3D chromatin folding. It shows how DNA sequence and folding patterns shape stochastic genomic processes in a crowded nucleus. In this paradigm, DNA-binding motifs primarily increase the local concentration of effector proteins, uncoupling protein binding to DNA from downstream processing of the genomic substrate. The replication and transcription factors that are confined in genomic pockets cycle ON and OFF chromatin but are prevented from diffusing away. A pocket functions as an effective trap only when global motif density is matched or exceeded by global levels of cognate proteins, making motif distribution a direct determinant of local protein concentration. Just as origins in a pocket share a common pre-RC pool, promoters - and enhancers in larger genomes-are “serviced” by a shared set of RNAPol2 complexes and transcription factors, which would explain why neighboring genes often display similar expression levels across organisms ^36^.

The activation of thousands of origins with relatively slow forks in each cell suggests that replicating chromatinized, actively transcribing DNA is intrinsically inefficient. Encounters with the transcription machinery and the topological strain around forks increase stalling and the risk of DNA-damage. Our results indicate that efficient replication in these conditions requires near-simultaneous activation of thousands of closely spaced origins, breaking the genome into small segments that individual forks can manage. If one fork stalls, another is typically within ∼1 kb to rescue it.

Given the high conservation of both chromatin architecture and the replication machinery, replication in other eukaryotes likely proceeds in a similar manner as in yeast. Even though individual peaks were not systematically analyzed, recent single-read replication profiles in human and Drosophila cells show EdU distributions consistent with dense origin arrays ^37,38^. Because the yeast genome is particularly gene-rich, it remains important to determine whether long, rarely transcribed intergenic regions in larger genomes are replicated in the same way or whether tightly packed origin arrays are mainly characteristic of the replication of transcriptionally active domains.

The long- and short-range cooperativity of origin activation (respectively, ≥10kbp and 0.5-2kbp, **Figs. 3D**, 6A–C) and the stimulation of fork velocities by nearby active origins (**Fig. 6E**) are likely the most efficient ways to overcome the topological constraints that hinder replication of long chromatin fibers. The slow firing of neighboring origins early in S phase - when fork density is low - suggests that the main barriers to fork progression are not collisions between forks and the transcription machinery but the inability of isolated Mcm2-7 helicases to efficiently unwind chromatinized DNA. Our analysis indicates that long chromatin fibers can be unwound efficiently only when many sites initiate nearly simultaneously along the chromosome. In light of our results, rapid activation of thousands of origins and efficient resolution of the resulting termination events should prove to be essential for optimal replication while mechanisms (if there are any) that facilitate the advancement of individual forks through long transcribed regions should prove to be accessory at best.

We can now finally explain why apparent changes in the replication program often accompany genome instability ^39,40^. Because most loci fire nearly simultaneously in the last quarter of S phase, replication timing itself is unlikely to preserve genome integrity. Apparent timing shifts observed under replication stress reflect instead the increased detectability of late replication, caused by a global decrease in EdU or BrdU consumption. We propose that replication stress reduces the number of early origin firing events per cell, which decreases nucleotide analog usage in early S-phase, and increases the labeling of late initiation events that are typically missed in normal growth conditions.

Thus, genome instability arises from a global drop in origin activation, not altered replication timing. Fewer firing events slow down replication, leave a significant number of genomic regions under-replicated, and increase the risk of chromosome abnormalities if cells enter mitosis. Any condition that causes the under-licensing of origins, such as reduced global levels of pre-RC components or mutations impairing pre-RC activation, DNA binding, or helicase function ^23,41^, will decrease origin firing and produce an altered timing profile in experimental conditions with limiting cellular nucleotide analogs. The apparent changes in replication timing are therefore symptoms of the true cause of genome instability: insufficient origin firing.

The idea that genome instability stems primarily from defective origin-firing kinetics reshapes how we interpret replication stress and the altered replication dynamics of cancer cells. The next challenge is to reassess replication behavior in individual cancer cells using NanoRep and the analytical framework developed here. Single-cell, single-molecule measurements will be crucial for determining how cancer-associated changes in origin activation drive genome instability.

## Acknowledgments

We thank the “Laverie” and Yeast facility (IGMM –CRBM) for yeast media preparations.

## Author contributions

LTN performed the EdU pulse and T chase replicate 1 and replicate 2 experiments and constructed the NTL3 strain. CJ-B performed the EdU pulse experiments (replicates 1 and 2). JG performed the EdU pulse and T chase replicate 3 experiment. BA and DR performed the machine learning modeling. MRL designed the experiments, analyzed the data, and wrote the manuscript.

## Data, code, and materials availability

ONT (Oxford Nanopore Technology) datasets with basecalled fast5 and bam files were submitted to the NCBI BioProject database (accession: BioProject ID PRJNA1474295, http://www.ncbi.nlm.nih.gov/bioproject/1474295). Raw pod5 files from the EdU pulse T chase rep 3 experiment are available upon request (NCBI BioProject does not accept pod5 files without basecalling). Perl scripts used for EdU/T peak calling and fv and t0n calculations are available in the NanoRep_pipeline.tar.gz file in Supplementary Materials.

