## Supplementary Materials for "Origins Anywhere and Everywhere: Genome replication requires cooperative activation of thousands of origins in each cell"

**The PDF file includes:**

Materials and Methods
Supplementary Text (Computational Methods)
Figures S1 to S11
References

**Other Supplementary Materials for this manuscript include the following:**

Data S1 to S3
Code files

### Materials and Methods

#### Yeast Strains

All strains have a W303 background and contain 5 copies of the hENT1 nucleoside transporter and seven copies of the Thymidine Kinase from Herpes Virus, both necessary for processing EdU added to the media. **RZ71** (*MATa ade2-1 his3-11,15 leu2-3,112 trp1-1 ura3-1 can1-100 GAL psi+ RAD5+ Rpb3-HA::TRP URA3::GDP-TK(7x) AUR1c::ADH-hENT1(5x) Δbar1::His3*) (*1*) was used in replicate 1 of the EdU pulse and T chase and the EdU pulse experiments (see **Fig. S2**). **NTL3** (*MATa trp1-1 can1-1000 leu2-3,112 his3-11,15 GAL psi+ RAD5+ ura3::URA3/GPD-TK(7x) AUR1c::ADH-hENT1(5x) bar1D::KANR ade2-1::promotorSIR3-NLS-EcoGII::ADE2*) (this study) was used in replicate 2 of the EdU pulse and T chase experiment. **PV1** (*MATa ade2-1 trp1-1 can1-1000 leu2-3,112 his3-11,15 GAL psi+ RAD5+ ura3::URA3/GPD-TK(7x) AUR1c::ADH-hENT1(5x) bar1D::KANR*) (*2*) was used in replicate 3 of the EdU pulse and T chase experiment.

#### NanoRep cell cultures and genomic DNA preparation for nanopore sequencing

EdU pulse Thymidine chase experiments

For replicates 1 and 2, cells were grown overnight at 30°C in SCD-URA to $OD_{600}$ ~0.3. The culture (200ml per time point) was synchronized with the addition of 0.15 µg/ml α factor for 3h30min at 30°C. Cells (OD~0.6) were pelleted and released from arrest into preheated (30°C) SCD-URA + 20 µM EdU (Carbosynth)+20µg/ml pronase (SIGMA). At 15, 25, and 35 min after release, cultures were mixed with 1x culture volume of preheated SCD-URA containing 2000x (over EdU) of Thymidine (SIGMA) (1000x for rep 2), for a final concentration of 1000x Thymidine (500x for rep.2), i.e. 20mM (10mM for rep. 2). Cells were then grown up to 60min after release, pelleted, flash frozen in liquid nitrogen and kept at -80°C until further processing. For replicate 3 cells were grown as above with the following modifications: YPD was used for the media, the OD before α factor addition was ~0.5, and the culture volume was 100ml per time point. There was also an additional 60min time point without any Thymidine addition.

EdU pulse experiments:

Cells were grown overnight at 30°C in YPD to $OD_{600}$ ~1. The culture (200ml per time point) was synchronized with the addition of 0.15 µg/ml α factor for 3h30min at 30°C. Cells were pelleted and released from arrest into preheated YPD +20µg/ml pronase (SIGMA). EdU was added to a final concentration of 25 µM at indicated times, as illustrated in Fig. S2B. All time points were collected 60min after release, pelleted, flash frozen in liquid nitrogen and kept at -70°C until further processing.

Spheroplasting and genomic DNA extraction:

Frozen cell pellets were re-suspended in Buffer Z (1 M Sorbitol +50mM Tris-HCl pH 7.5) at ~$10^9$ cells/100µl, thawed on ice and incubated with 25 U of Zymolyase (AMSBIO) + βME (10mM final), for 30 min at 30°C (no shaking or 10" at 300rpm every 10min to minimize DNA breakage).

High Molecular Weight DNA was then extracted from pelleted spheroplasts with the MagAttract HMW DNA (Qiagen), or the Nanobind CBB Big DNA (Circulomics) kits (EdU pulse chase, rep. 1, 2 and 3). Replicates 1 and 2 were treated with an additional 0.4x MagNa beads (Sera-Mag Magnetic Speed-beads (FisherSci, cat.#: 09-981-123))(*3*) purification step after extraction to deplete shorter fragments and reduce RNA contamination. Genomic DNA from EdU pulse replicates 1 and 2 was purified with the Monarch HMW DNA extraction kit for cells and blood (NEB), according to the manufacturer's protocol. An additional optional RNAse A degradation step for EdU pulse chase rep. 3 (1mg/ml, 1hr to o/n at 37°C, no shaking) was used when the ratio of DNA levels measured by nanodrop (measures DNA and RNA) and

DNA levels measured by Qubit (measures dsDNA only) exceeded 2. Yeast genomic DNA preps are often heavily contaminated with RNA despite the RNAse step in the kit purification protocols, and since excess RNA inhibits library preparation it is very important to minimize RNA contamination.

Nanopore sequencing:

200ng to 400ng from each genomic DNA sample (measured with the Qubit ds DNA kit) were used for nanopore sequencing library preparation with the Rapid Barcoding Kit (Oxford Nanopore), (SQK-RBK004 for 9.4.1 flow cells or V114.24 for 10.4.1 flow cells), according to the manufacturer protocol.

Libraries prepared with the Rapid Barcoding Kit were mixed, cleaned, and concentrated with 1x MagNA or Ampure beads as described in the Rapid Barcoding Kit protocol.

The library mix was loaded on a R9.4.1 Flow cell (Oxford Nanopore) (EdU pulse chase, rep. 1 and 2 and EdU pulse experiments) or a R10.4.1 Flow cell (EdU pulse chase rep 3) and sequenced with the Minion device (Oxford Nanopre) for 48 to 72 hrs without the real time base calling option.

**Nanopore sequencing data analysis**

For replicates 1 and 2 of the EdU pulse and T chase experiment and for the EdU pulse experiment base calling was performed on raw fast5 files, using guppy basecaller (ONT, Community - Downloads (nanoporetech.com)) with res_dna_r941_min_modbases-all-context_v001.cfg. Fastq files were then extracted from base-called fast5 files using GitHub - nanoporetech/ont_fast5_api: Oxford Nanopore Technologies fast5 API software). Demultiplexing and genome alignments were done with the guppy barcoder and the guppy aligner, respectively (*S.cerevisiae* reference genome: S288C_reference_sequence_R64-3-1_20210421.fasta). EdU detection was performed with DNAscent 3.1.2 (https://github.com/MBoemo/DNAscent/releases) (*4*). Thymidine bases with an EdU probability of at least 0.85 were considered as positive EdU calls.

For replicate 3 of the EdU pulse and T chase experiment, base calling was performed on raw pod5 files, using dorado (0.9.0) basecaller (ONT, Community - Downloads (nanoporetech.com)) with the hac configuration. Fastq files were then extracted from base-called untrimmed bam files using samtools. Demultiplexing was done with the guppy barcoder on the resulting fastq files. Guppy barcoder can still be used with the Rapid Barcoding V114.24 kit provided that only the first 12 barcodes are used in library preparation, as the first 12 barcodes are the same in the older and newer versions of the kit. We used guppy barcoder instead of dorado demux because guppy barcoder emits a barcoding summary with barcode alignment scores (a feature that was eliminated from the dorado demux barcoding summary), which enables more customizable and less stringent barcode annotations: we retain all barcode annotations with a score of at least 40. The genome alignment was done using dorado aligner (*S.cerevisiae* reference genome: S288C_reference_sequence_R64-3-1_20210421.fasta) on adapter trimmed base called bam files. EdU detection was performed with DNAscent 4.0.3 (https://github.com/MBoemo/DNAscent/releases) (*4*). Thymidine bases with an EdU probability of at least 0.75 were considered as positive EdU calls.

EdU/T peak detection:

Perl scripts used for the analysis described below are available in NanoRep_pipeline.tar. The order in which the scripts are executed is listed in pipelinelist.txt included in the tar file. We first determined the number of EdU bases per total number of Thymidine bases per 400bp bin of each read. Watson and Crick reads were treated separately for the entire analysis pipeline. We then isolated all the reads that had at least one 400bp bin with incorporated EdU, and identified all full peaks on each one of those reads.

Full peaks are defined as regions of at least three consecutive 400bp bins (in a 5' to 3' direction of the reference Watson strand) with non-zero EdU/T. The first bin of a full peak (i.e. the upstream edge) has the minimum EdU/T ($(EdU/T)_{upedge}$) of all the bins upstream of the bin with the highest EdU/T (the peak,) and the last bin (downstream edge) has the minimum EdU/T ($(EdU/T)_{downedge}$) of all the bins downstream of the peak.

We then normalized EdU/T values for each 400bp bin in each full peak with the maximum $(EdU/T)_{peak}$ per time point, and calculated the timing of origin firing ($t_{0n}$) and the upstream and downstream fork velocity for each full peak as respectively described in **Figs. 3A and 4A**. Only full peaks with $[(EdU/T)_{peak}–(EdU/T)_{edge}]/ (EdU/T)_{peak} \geq 0.2$ were used for $t_{0n}$ and fork velocity calculations, to deplete false positive peaks in which EdU/T differences are due to random fluctuations in EdU incorporation that are not linked to rapid cellular EdU usage during replication.

**Supplementary Text: Computational methods**

### S1 Predicting the origin firing efficiency solely from the DNA sequence immediately surrounding mapped origins

#### Materials and Methods

#### Datasets

Two experimental datasets provided the efficiency targets. The **NChAP 18 min dataset** ($n$ = 4,750 origin (ARS) windows) reports normalized NChAP peak height at 18 min post-release from G1normalized as described in (*1*) (Figs. S1A and 7A). The NanoRep EdU pulse T chase rep1. dataset ($n = 2{,}654$ windows) reports origin peaks mapped from average EdU/(T*n_reads) per 400bp locus across the cell population (Fig.1A) obtained from single-molecule NanoRep measurements at 25 min after release from G1 (Fig. S2). Both datasets represent 500 bp windows (±250 bp from each ACS coordinate). Efficiency values are right-skewed; a log1+$x$ transformation was applied before computing losses and metrics.

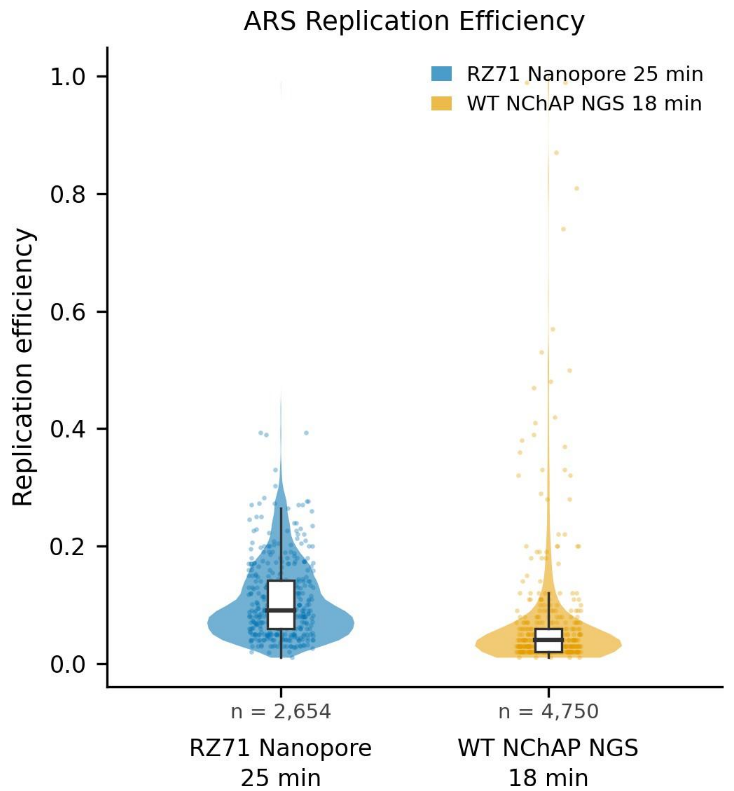


**Figure 1: Distribution of ARS replication efficiency for both datasets.** Both distributions are heavily right-skewed, motivating the log1+$x$ transformation applied prior to model training.(*5*)

#### Sequence Representations

The following sequence representations were evaluated as model inputs:

**K-mer frequency vector** (SVM, RF, CNN) : overlapping k-mer counts for $k \in [2, 3, 4, 5]$ converted to relative frequencies, giving 1,360 values per sequence. Encodes nucleotide composition without positional information.

**Integer encoding** (Triplet Fusion CNN): nucleotides stored as integer indices (0–3 for A/C/G/T; 4 for N); the network converts these to one-hot internally during the forward pass.

**DNABERT-2 BPE tokenisation** (DNABERT-2): the sequence is split into variable-length tokens using a byte-pair encoding (BPE) tokeniser pre-trained on large DNA corpora. BPE iteratively merges the most frequent character pairs into longer vocabulary tokens, so common nucleotide strings are represented as single units. Each token is mapped to a 768-dimensional embedding with an added positional embedding, giving a token-sequence matrix as input to the transformer.

### Models

**SVM.** Trained on the k-mer frequency vector (standardised). Four kernels (linear, RBF, polynomial, sigmoid) were compared via GroupKFold(5) CV with fixed hyperparameters ($C$ = 1 for linear/poly/sigmoid, $C$ = 10 for RBF; $\gamma$ = scale).
**Random Forest.** Ensemble of 500 trees on the k-mer frequency vector.
**K-mer frequency CNN.** Four parallel 1D convolutional branches with filter widths $k \in [5, 11, 21, 41]$, global max-pooling, then a shared FC head (256 → 128 → 1, dropout $p$ = 0.4; ∼38,000 parameters).
**Triplet CNN.** Three consecutive ARS windows (left neighbour, target, right neighbour) each encoded by a shared convolutional stem (kernel 7, 32 filters, average pooling); concatenated and decoded by a FC head (96 → 64 → 1; 7,201 parameters). Pairs separated by >10 kb were excluded, yielding 2,016 triplets.
**DNABERT-2.** Frozen DNABERT-2(5) (117M parameters) produced per-token embeddings (dim 768); a trainable head (896,769 parameters) — residual CNN, 8-head attention pooling, MLP 256 → 128 → 64 → 1 — decoded them to a scalar.

### Training and Evaluation

Deep learning models were trained with Adam ($\eta$ = $10^{-3}$, weight decay $10^{-4}$), ReduceLROnPlateau (factor 0.5, patience 12, min $\eta = 10^{-5}$), early stopping (patience 30 epochs on validation MSE in log space), and gradient clipping ($\ell_2 \leq 1.0$). Loss was MSE on log1+$x$ targets; predictions were back-transformed before metric computation. All models were implemented in PyTorch.
All models were evaluated by 5-fold GroupKFold cross-validation. Groups were defined by overlap-cluster (ACS coordinates within 500 bp on the same chromosome placed in the same cluster). Reported metrics — $R^2$ , RMSE and Spearman $\rho$ — were computed on held-out fold predictions assembled across all five folds.

### Results

Cross-validated predictions (GroupKFold-5) were assembled from held-out folds and evaluated in raw space (Table 1). Spearman $\rho$ is the primary metric given the heavy-tailed efficiency distribution. Across both datasets, none of the models could make a strong prediction. All $R^2$ values are negative across both datasets and all models, indicating that no model captures systematic variance in replication efficiency beyond the global mean.

### Conclusion

The low Spearman correlations across all models indicate that raw DNA sequence composition carries only a weak, statistically detectable signal for ARS replication efficiency in *S. cerevisiae*. These results imply that replication timing at the population level is largely determined by factors beyond primary sequence immediately surrounding (within 250bp upstream or downstream) a mapped origin.

Table 1: Cross-validated performance ($R^2$ raw space; Spearman $\rho$) on both datasets.

| | **WT NChAP 18 min** | | **RZ71 Nanopore 25 min** | |
|---|---|---|---|---|
| Model | $R^2$ | Spearman $\rho$ | $R^2$ | Spearman $\rho$ |
| SVM | −0.044 | 0.061 | −0.094 | 0.055 |
| Random Forest | −0.050 | 0.075 | −0.077 | −0.031 |
| XGBoost | −0.051 | 0.040 | −0.121 | 0.005 |
| DNABERT-2 | −0.051 | 0.083 | −0.004 | 0.078 |
| Triplet CNN | −0.046 | 0.169 | −0.068 | 0.098 |

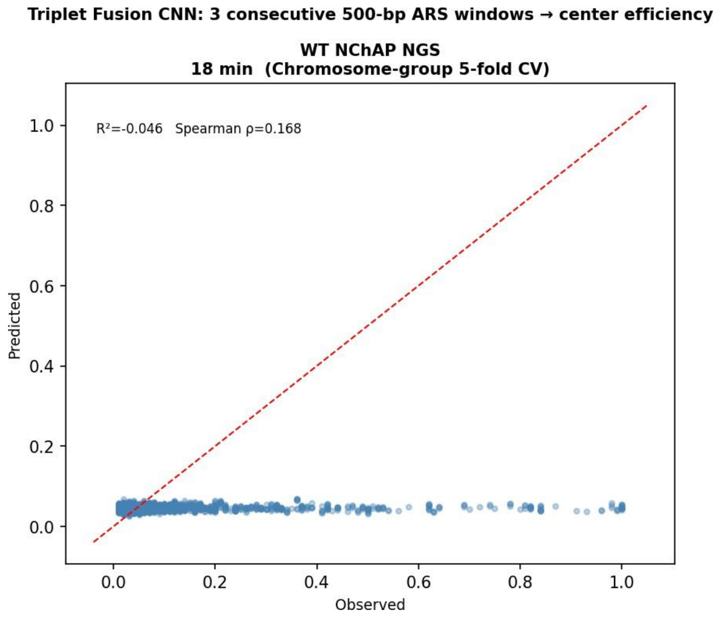


Figure 2: Triplet CNN observed vs. predicted on WT NChAP 18 min ($\rho$ = 0.169).

**S2 Modeling of Genome wide Origin Firing Probabilities (the corrected ACS model)**

**Data sources.** ACS motif density was obtained from the count of all possible ACS motifs (Fig. S1A) per 400 bp non-overlapping contiguous genomic bin across the *S. cerevisiae* genome.
ORC peak calls (nOrcPeaks) are from (*6*) and NanoRep firing probabilities for time windows 0–6 min and 6–8 min are from this study (Data S3, Fig. 7A).
Hi-C contact matrices for *S. cerevisiae* S-phase wild-type were obtained from (*7*) (three-replicate average, GSE309730) at 1 kb resolution, ICE-balanced. A second asynchronous wild-type dataset ((*8*), GSE227687) was used for comparison.
Sir3 occupancy from Sir3EcoG2 *in vivo* footprinting data (meA/(A·readn), Data S3) was obtained from (*9*); raw values were clipped at zero and min–max normalized to 0, 1 per genome.
RNAPol2 ChIP signal (Rpb3-HA, G1 phase) was obtained from (*1*) . It was interpolated linearly within each chromosome for missing bins, and min–max normalized to 0, 1. Both the ChIP and *in vivo* foot printing signals were additionally smoothed per chromosome with a uniform moving average of 20 kb (50 bins, mode="reflect").
Three-dimensional coordinates (nm) for each genomic locus were obtained from the 3D model of the *S. cerevisiae* genome in (*10*).

<u>A mechanistic model of firing probabilities</u>

We computationally analysed the correlation between the information in these datasets with the origin firing probability measured by NanoRep (see Figs. S6 and S7). To do so, we tested the Pearson correlation of the following data with the 0-6 ($1^{st}$ half of S-phase) and 6-8 ($3^{rd}$ quarter of S-phase) NanoRep firing probabilities (see Fig. 7A and Data S3).

**Baseline ACS density.** Raw ACS counts (n_acs per 400 bp) were smoothed per chromosome with a 10 kb uniform moving average (25 bins, mode="reflect") to produce the baseline ACS_10kb used in all subsequent corrections and identified in Fig. 5B-C and S4 to directly correlate with the probability of firing early in S-phase.

**ORC proximity to Hi-C.** ORC-containing 400 bp bins were mapped to the 1 kb Hi-C grid. For each ORC bin, ICE-balanced contact counts were extracted from the cooler matrix. For every non-ORC bin *i*, the ORC proximity score p_ORC(*i*) was the sum of balanced Hi-C contacts with all ORC-containing bins. Scores were converted to percentile ranks within each chromosome and scaled to 0, 1.

**3D centromere distance.** Three-dimensional coordinates (nm) for each genomic locus were obtained from a 3D model of the S. cerevisiae genome(*10*). For each 400 bp bin, the Euclidean distance d_3D(i) to the centromere of its chromosome was computed. The centromere boost factor was: f_cen(i) = 1 + b exp(-d_3D(i) / c)
with b = 50 and decay constant c = 50 nm, matching the parameterization used in prior simulation work.

**Telomere decay.** The linear distance from each bin to the nearest chromosome end was computed as d_telo(i) = min(bp, chr_len − bp) / 1000 (in kb). The telomere suppression factor was: f_telo(i) = 1 - A exp(-d_telo(i) / λ)
with A = 0.87 and λ = 14 kb, optimized to minimize residual distance correlation in the corrected ACS.

**Sir3 suppression.** The smoothed (20 kb) and normalized Sir3 occupancy signal s_20(i) was used as a proxy for Sir complex occupancy and its potential effect on origin firing delay at chromosome ends shown in Fig. S5. The suppression factor was:

$$f\_sir3(i) = 1 - \alpha\, s\_20(i)$$

with $\alpha = 10$. The observed correlation between telomere decay and Sir3 suppression is consistent with our hypothesis that the Sir complex is responsible for the delay in origin firing at chromosome ends.

**RNAPol2 boost.** The smoothed (20 kb) and normalized RNAPol2 ChIP signal p_20(i) was used as a proxy for transcription-associated chromatin accessibility. The boost factor was:

$$f\_pol2(i) = 1 + \beta\, p\_20(i)$$

with $\beta = 2$ (smoothed version). Only the smoothed, low-gamma variant was retained as the best-performing parameterization.

**Multiplicative correction.** The fully corrected ACS density was computed as the product of all independent factors:

```
acs_corrected = ACS_10kb
        × (1 + 50·exp(-d3d_cen / 50))     ← centromere 3D boost
        × (1 + 20·hic_prox)               ← ORC proximity boost
        × (1 − 0.87·exp(-d_telo / 14))    ← telomere decay
        × (1 + 2·pol2_sm20)               ← RNAPol2 boost
```

The final corrected ACS density at each 400 bp genomic bin is computed as a product of four independent multiplicative factors applied to ACS density (smoothed at 10 kb). First, a centromere 3D boost $(1 + 50\cdot e^{(-d3d/50)})$ captures the spatial advantage of loci close to the centromere in the Duan et al. 3D model, where d3d is the Euclidean distance in nm. Second, an ORC proximity boost $(1 + 20\cdot prox)$ reflects the summed Hi-C contact count between each bin and all ORC ChIP-seq-positive bins. Third, a telomere decay function $(1 - 0.87\cdot e^{(-d\_telo/14)})$ simulates the Sir3 occupancy signal that correlates with the suppression of early origin firing near chromosome ends (Fig. S5), where d_telo is the linear distance to the nearest telomere in kb. Fourth, a RNAPol2 boost $(1 + 2\cdot pol2\_20kb)$ adds a modest up weight at transcribed regions using the RNA Polymerase II ChIP-seq signal smoothed at 20kbp. The four factors are multiplied together with ACS_10kbp to produce the final corrected ACS value at each bin.

**Correlation analysis.** Pearson correlation coefficients were computed between each feature (or corrected ACS variant) and each NanoRep firing probability target across all autosomal bins (chromosomes 1–16). Only the best-performing variant per model family was retained for presentation in Fig. S6. Spatial ACS and ORC features (Hi-C contact-weighted averages of ACS density or ORC peak count in 3D neighborhoods) were computed via sparse matrix multiplication of the Hi-C contact matrix with the smoothing -aggregated signal using contact-type filtering and row-normalized averaging. Sir3 and RNAPol2 multiplicative variants (ACS × Sir3, ACS × Pol2) used the same normalized signals (with smoothing).

**S3 Predicting origin firing probabilities with machine learning (Fig. S8-S11)**

**Prediction models.** Firing probability was predicted at 4 kb resolution (average of ~10 adjacent 400 bp bins per feature) using Ridge regression (α = 1.0), Random Forest (max_depth = 10, 150 trees), and XGBoost (max_depth = 4, 150 trees, learning_rate = 0.1). Features comprised the 14-dimensional vector: ACS_10kbp, ORC peak count, Sir3 occupancy (raw and 20 kbp smoothed), RNAPol2 signal (raw and 20 kbp smoothed), telomere distance, 3D centromere distance, ORC proximity, and spatial ACS/ORC proximity (two Hi-C datasets each). Models were evaluated via leave-one-chromosome-out cross-validation (16 folds, one chromosome held out per fold). Reported metrics are Pearson r between predicted and observed values on held-out chromosomes. All 12 features are computed per 400 bp bin and aggregated to 4 kb by averaging within each 4 kb window. Features are z-score standardized for Ridge; tree models use raw values.

**Software.** Analyses were performed in Python using scikit-learn, XGBoost, SciPy, NumPy, Pandas, and Matplotlib. Hi-C data were processed via Cooler. The full analysis code is available in the data_analysis/ directory of the repository.

**Feature description:** the following text contains the description of the 12 features used in the machine learning models

**1. ACS density (10 kbp) — `acs_10kb`**

Number of ACS (ARS consensus sequence) matches per 400 bp bin, smoothed with a uniform filter (`scipy.ndimage.uniform_filter1d`, mode="reflect") at a bandwidth of 10 kb (25 bins), then averaged to 4 kb.

<u>What it captures:</u> The local density of potential origin recognition sequences. This is the baseline linear feature. It measures how many ARS consensus elements are present near each bin, without any 3D information. This is the same feature identified in Figs. 5B-C and S4A-B, as having a genome –wide direct average correlation with origin firing early in S-phase (first 75% of S-phase (0-8min from the start)). The 10 kb smoothing removes bin-to-bin noise while preserving regional variation.

<u>Why 10 kb?</u> Prior analysis showed that 10 kb is the optimal smoothing scale for ACS density against NanoRep firing probability, balancing noise reduction against retention of origin-scale signal. As shown in Fig. 5C (middle panel), 10kbp is the smallest genome bin, for which we identified an early origin firing advantage at the high end of the local ACS density distribution: 5kbps bins with a local average ACS density from 3 to 7 ACS/kbp, do not fire on average any earlier than 10kbp bins with a local average ACS density ranging from 2.5 to 4.5 ACS/kbps.

**2. ORC peaks (20 kbp) — `n_orc_peaks`**

Integer count of ORC (origin recognition complex) ChIP-seq peaks per 400 bp bin, smoothed with a uniform filter at 20 kb (50 bins). The smoothed value replaces the raw count for all downstream uses: the ORC proximity flag (features 7–8), the ORC 3D density (features 10, 12), and this linear feature itself. Note that ChIP-seq is not very sensitive nor very quantitative and only detects the sites with moderate to high ORC occupancy, which by definition have a higher probability of firing early.

What it captures: The presence and density of experimentally mapped ORC-binding sites. Unlike the raw binary peak presence, the 20 kb smoothed version produces a continuous score that reflects the broader ORC-binding landscape. The 20 kb bandwidth was chosen because ORC peaks are sparse (~300–400 genome-wide), and smoothing spreads their influence to adjacent bins that may be functionally linked through a shared local chromatin environment. The smoothed value is used as a threshold (`> 0`) to flag Hi-C bins as "ORC-bound" for the 3D proximity computation, ensuring that isolated single-bin peaks do not dominate the Hi-C contact aggregation.

**3. Telomere distance — `d_telo`**

Linear distance from each bin to the nearest chromosome end, in kilobases: `min(bp, chr_length - bp) / 1000`.

What it captures: As shown previously (*1*) and confirmed in this study, the left and right subtelomeric domains (0–25 kb from each end) are on average among the latest-firing regions on each chromosome (Figs 7A, S4 and S5). This feature provides a simpler linear proxy for this telomere positional effect than Sir3 occupancy levels, which are probably the cause of the observed delay in origin firing at chromosome ends (see Fig. S5).

**4. 3D cen. distance — `d3d_cen`**

Euclidean distance (nm) from each bin to the centromere in the 3D chromosome model from (*10*).

What it captures: 3D proximity to the centromere. In the yeast nucleus, centromeres cluster near the spindle pole body, creating a radial organization where centromere-proximal loci are spatially constrained and typically replicate early. The Duan model provides (x, y, z) coordinates for ~9,000 loci per chromosome from a population-averaged Hi-C-based 3D reconstruction. This feature captures the centromere-proximal replication advantage that is not fully explained by linear distance alone (e.g., loci far from the centromere in linear sequence can be spatially close to it in 3D).

**5. Sir3 occupancy (20 kbp) — `sir3_norm_20kb`**

Sir3 population averaged occupancy (from Sir3EcoG2 in vivo foot printing,(*9*), Fig. S5, Data S3) clipped to non-negative values, normalized to the [0, 1] range per chromosome, and smoothed with a uniform filter at 20 kbp.

What it captures: Sir3 is a core component of the SIR (Silent Information Regulator) complex that establishes and maintains heterochromatin at the HML and HMR silent mating-type loci and subtelomeric regions (see Fig.S5, Data S3) (*9, 11*). Subtelomeric regions with high levels of Sir3 typically fire in the last S-phase quarter (Figs. 7A and S5, Data S3). The 20 kbp smoothing matches the scale of Sir3-spread domains (typically 3–20 kb from telomeres).

**6. RNAPol2 ChIP (20 kbp) — `pol2_norm_20kb`**

RNAPol2 (Rpb3 subunit, G1 phase) ChIP-seq signal from (*1*), normalized to [0, 1], gaps interpolated per chromosome, and smoothed at 20 kbp.

What it captures: Active transcription — RNAPol2 enrichment marks euchromatic, actively transcribed regions, and correlates with gene expression (mRNA) levels. Genome wide RNAPol2 from G1 was used to assess if transcription levels influence origin licensing, which happens in G1. The 20 kb smoothing matches the typical size of Replication Domains identified in this study.

**7. 3D ORC proximity (D'Asaro2025) — `hic_prox_das2025_sphase_wt`**
and
**8. 3D ORC proximity (GSE227687) — `hic_prox_gse227687_wt`**

**Datasets:**
D'Asaro2025 (early S-phase) GEO: GSE309730(*7*): Three biological replicates of early S-phase *S. cerevisiae* (20 minutes after α-factor release from G1 arrest), sequenced to ~100 million contacts per replicate. Contacts are aggregated at 1 kb resolution and ICE-balanced. The three replicates are averaged to reduce noise.

Dataset GSE227687 (asynchronous WT) GEO: GSE227687 (*8*): Asynchronous wild-type *S. cerevisiae*, 1 kb resolution `.mcool`, ICE-balanced. This dataset captures the population-average contact map without cell-cycle synchronization, providing a broader but less S-phase-focused view of 3D organization.

Feature construction: For each 400 bp bin, the sum of ICE-balanced Hi-C contact counts to all 1 kb bins that overlap an ORC peak (smoothed `n_orc_peaks > 0` at 20 kb), converted to a per-chromosome percentile rank [0, 1]. The raw proximity score is: "how much Hi-C contact does this bin have with any ORC-bound locus, anywhere in the genome (intra- and inter-chromosomal)?" ICE normalization corrects for technical biases (GC content, mappability, fragment length). The percentile rank makes the score comparable across chromosomes.

What it captures: A bin that is frequently in 3D contact with ORC-bound loci receives a high proximity score — even if those loci are separated by megabases of linear sequence or lie on different chromosomes. This is fundamentally different from linear genomic distance. The biological rationale: origins that are spatially clustered near ORC-bound loci may benefit from a shared pool of limiting initiation factors (ORC, Cdc6, Cdt1, Mcm2–7), increasing their firing probability. This feature is used multiplicatively in the "ACS × 3D ORC proximity" bar chart entry (factor `1 + 20 × proximity`).

**9. ACS 3D density (D'Asaro2025) — `sp_acs_das2025_sphase_wt`**
and
**11. ACS 3D density (GSE227687) — `sp_acs_gse227687_wt`**

For each 1 kb Hi-C bin, the contact-weighted average ACS density (10 kb) of all other bins that contact it in 3D: $sp\_acs[i] = \Sigma_j(acs_j \cdot w_{ij}) / \Sigma_j w_{ij}$ where $w_{ij}$ is the ICE-balanced contact count from cooler pixels (intra-chromosomal contacts only; inter-chromosomal contacts are excluded to avoid combining unrelated chromosome territories). This is the 3D-neighbourhood average of ACS density, not a linear smooth.

What it captures: A bin that contacts ACS-dense regions in 3D receives a high "3D ACS density" even if its own local ACS density is low. Conversely, a bin that contacts ACS-poor regions has a low 3D ACS even if it lies in an ACS-dense linear neighbourhood. This replaces the implicit assumption of linear smoothness with the actual 3D contact neighbourhood measured by Hi-C. The difference between the linear ACS density (feature 1) and the ACS 3D density reveals how much of the spatial ACS signal is due to 3D contacts with distal ACS-rich regions versus local linear ACS density.

Why intra-only? Inter-chromosomal contacts in yeast are sparse and noisy; including them adds variance without improving the spatial ACS estimate. Previous systematic testing showed that intra-chromosomal contacts produce the strongest correlation with replication timing.

**10. ORC 3D density (D'Asaro2025)(*7*) — `sp_orc_das2025_sphase_wt`**
and
**12. ORC 3D density (GSE227687)(*8*) — `sp_orc_gse227687_wt`**

Same procedure as ACS 3D density (features 9, 11), but using the 20 kb smoothed ORC peak count as the value column instead of ACS density:

$$sp\_orc[i] = \Sigma_j(ORC[j] \cdot w_{ij}) / \Sigma_j w_{ij}$$

where `ORC[j]` is the 20 kbp smoothed ORC peak count at bin j.

What it captures: How much ORC signal reaches each bin through its 3D contact neighborhood. A bin may have zero ORC peaks locally but contact regions rich in ORC peaks in 3D space, receiving a high ORC 3D density. This is distinct from the ORC proximity score (features 7–8): the ORC proximity score is a binary-derived (ORC yes/no) contact sum converted to percentile rank, while the ORC 3D density is a continuous weighted average of the actual ORC peak count over the 3D neighborhood. The continuous version may capture more subtle variation in ORC dosage than the binary proximity flag.

**Software.** Analyses were performed in Python 3.12 using scikit-learn, XGBoost, SciPy, NumPy, Pandas, and Matplotlib. Hi-C data were processed via Cooler. The full analysis code is available in the data_analysis/ directory of the repository.

**Conclusion:**

Figs. S6 and S7 show that the features described above increase the Pearson correlation between the 10kbp smoothed chromosomal ACS density distribution and the 0-6min and 6-8min firing probabilities measured by NanoRep (Fig. 7A and Data S3) up to r=0.39. We also benchmarked this biologically principled multiplicative correction of ACS density by comparing it with three linear and nonlinear machine learning models that are trained in a leave-one-chromosome-out cross-validation to predict the same 0-6min and 6-8min firing probabilities (Fig S8). The results show that the nonlinear models do not have advantage over the simpler ridge regression, and that in general these predictors do not significantly outperform the multiplicative ACS correction ($r \approx 0.39$–$0.41$, Figs. S9-S11). With respect to these models, the interpretable nature of our multiplicative approach interestingly shows (Fig. S6) that, while the ORC linear density improves the correlation with respect to the ACS density alone, considering the 3D ORC density (using Hi-C contact maps) provides a further boost to the final prediction, in line with the definition of the ORC "trap" model that we propose in Fig. 7B.

Both the corrected ACS and the machine learning prediction models are fairly successful at localizing the broader genomic regions (~20-30kbps) that have a high probability of firing in the first three quarters of S-phase (Figs. S9-S10). They are less good, however, at predicting the correct (observed) firing probability and estimate the local variability in firing probabilities for regions that fire “early” i.e. that mostly fire in the first 75% of S-phase across the cell population and “late” i.e. that predominantly fire in the last S-phase quarter across the cell population. The centromeric 3D boost function in the corrected ACS model tends to overestimate the firing probability of regions in close proximity to centromeres in 3D space and underestimate the firing probability of early regions distant from centromeres. The machine learning prediction models tend to even out local variations in firing probabilities across each chromosome, respectively underestimating or overestimating the early firing probabilities of early and late regions. This is not surprising. Both modelling methods compute a firing probability function that represents the best average fit to the experimental data across the entire genome, which by definition “smooths” away any variability in origin firing probabilities caused by local specificities of the chromatin environment. In other words, in order to identify features and extract global trends that influence origin firing genome wide, either modeling approach has to compromise and “neglect” the local variability of examined features that might have a more localized effect on origin firing dynamics in a specific genomic region. Consequently, neither model accounts for the localized boosting effect of moderate Sir3 occupancy on early origin firing in chromosome bodies (Fig. S5), nor for the reasonable possibility that despite an average general positive effect, RNAPol2 occupancy might have a localized negative effect on early origin firing in certain conditions due to RNAPol2 competition with ORC for binding to the same chromatin substrate.

Furthermore, the models are handicapped by the low sensitivity and low resolution of ORC ChIP-seq, and Hi-C datasets, which can only reliably detect the strongest and most frequent ORC binding, and intra-chromosomal 3D contacts in the cell population. Consequently, moderate to low ORC occupancy and less frequent 3D chromosome conformations that may contribute to local origin firing variability are absent from the input datasets and do not contribute to either modeling approach.

### Supplementary Figures

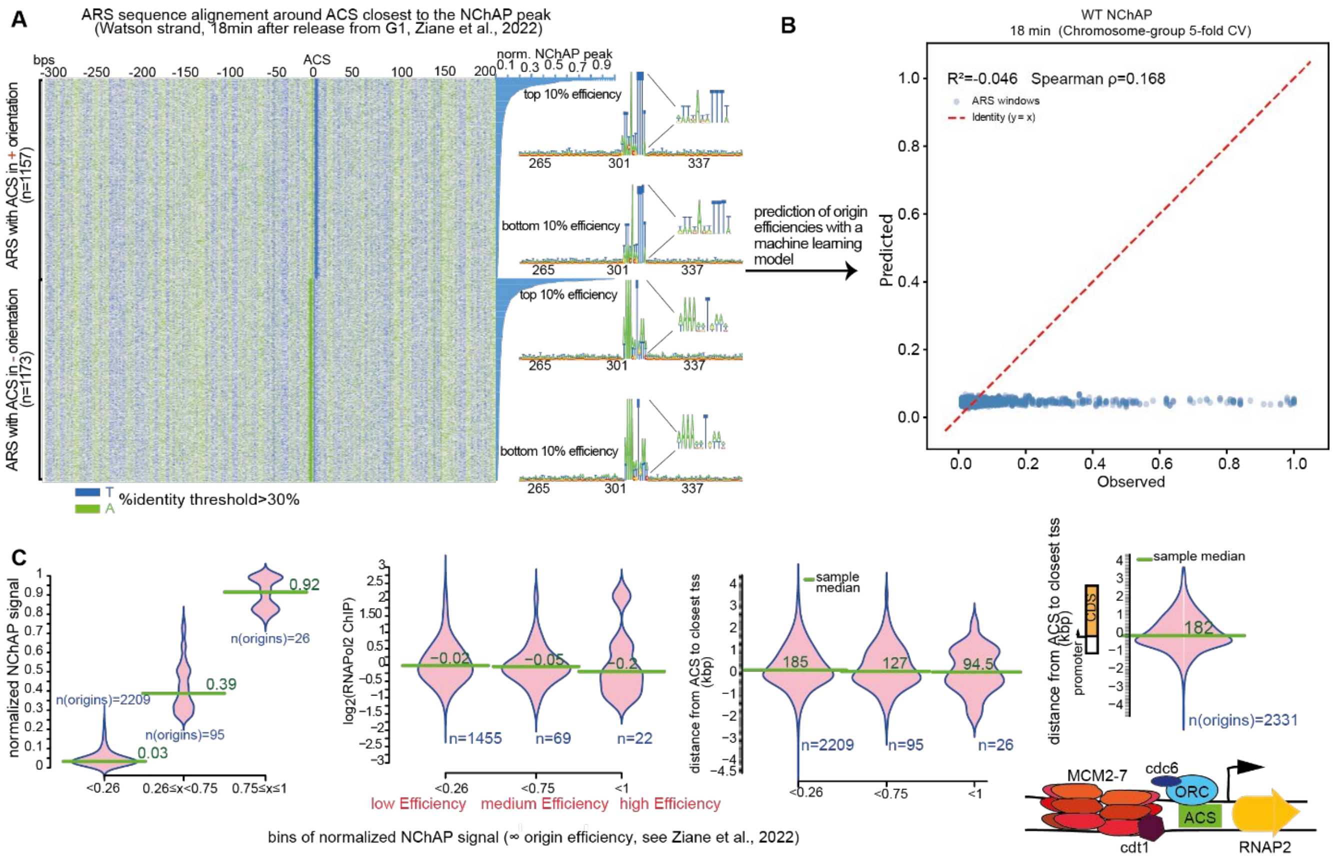


**Fig. S1. DNA sequence in the immediate vicinity of an origin does not determine origin efficiency as measured by population based NChAP (*1*). A.** Sequences of all Origins (ARS) mapped in (*1*) were centered around the ACS motif ((T|A|G)(T|A)(T|A)(T|C|A)A(T|C|A)(A|G|T)TTT(T|A)) closest to the center of the NChAP peak, and sorted by origin efficiency (the height of the NChAP peak). Origins with the central ACS in the + or – orientation were treated separately. Jalview was used to determine the % identity of aligned sequences. The consensus sequences from the top and bottom 10% of origins on the efficiency scale are shown on the right of the alignment. **B.** Triplet Fusion CNN: 3 consecutive 500-bp ARS windows → center efficiency. Observed versus predicted ARS replication efficiencies from 5-fold cross-validation of a triplet CNN using three consecutive ARS windows (left neighbor, target, and right neighbor) as input. Results are shown for 4,229 triplets (inter-window gap ≤ 10 kb). Each point represents a target ARS window. The dashed red line indicates the identity line (y = x). Spearman $\rho = 0.168$; $R^2 = -0.046$. **C.** Origins from A (+ and – ACS motifs were treated together) were sorted into efficiency quartiles (**first from the left**). The density distributions of RNAPol2 occupancy (midlog wt cells, data from (*12*)) (**second from left**) or distance from the tss of the gene closest to the central ACS (**third from the left**) were determined for each quartile of origin efficiency. The density distribution of the distance from the tss of the gene closest to the central ACS for all mapped origins from A is shown in the **rightmost** panel.

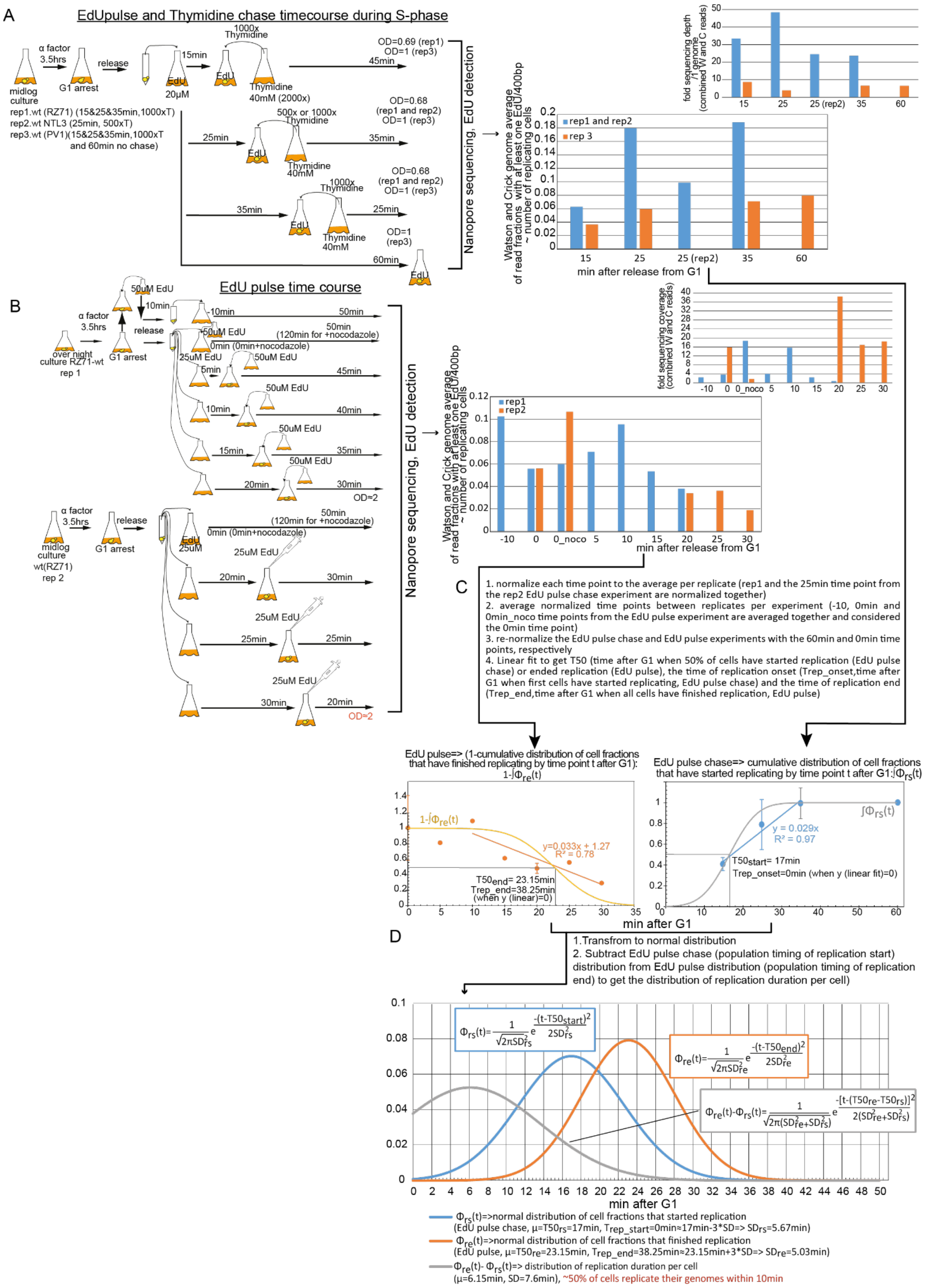


**Fig. S2. S-phase duration in single cells derived from EdU pulse, and EdU pulse and T chase time courses. A.** Diagram of the EdU pulse and T chase protocol (**left**). Bar graph showing the fraction of reads that have incorporated EdU out of all reads in each time point (**right**). The bar graph in the **top right corner** shows the average sequence coverage, i.e. the number of base pairs in all reads per time point divided by the size of the yeast genome in bps, which gives the average number of times the sequenced reads cover the yeast genome for each time point. **B.** Same as A. but for the EdU pulse experiment. **C.** the text on top describes the normalization procedure that transforms the raw data from B into cumulative distribution functions of cell fraction finishing replication (**bottom left**), and starting replication (**bottom right**). **D.** shows the normal distribution functions derived from the cumulative distribution functions from C.

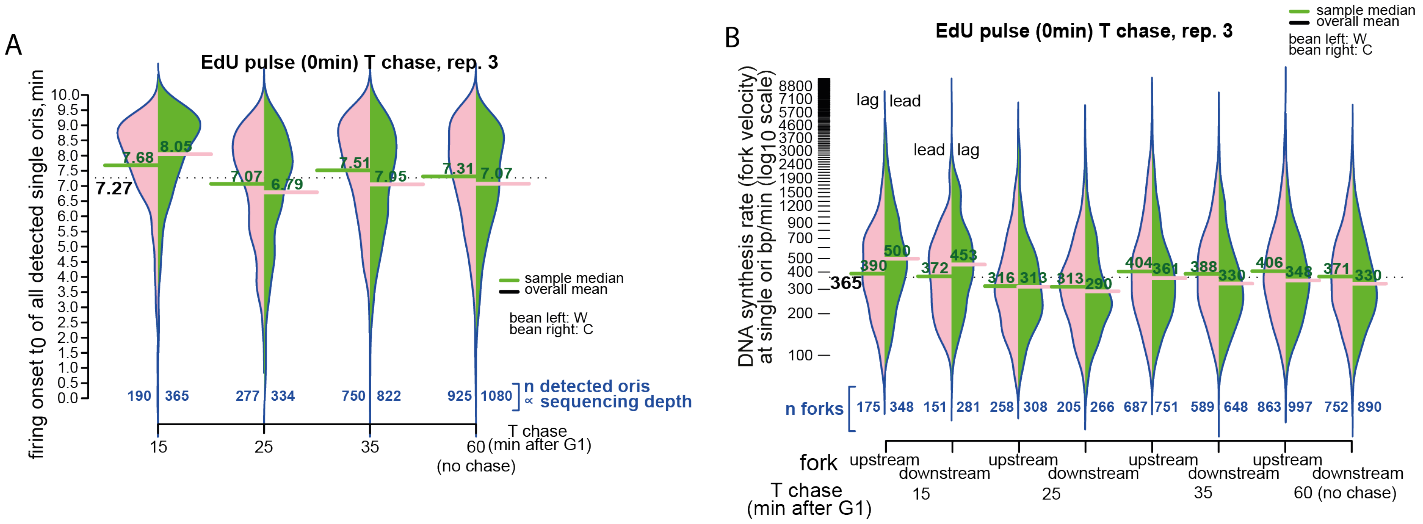


**Fig. S3. A-B.** like Figures 3B and 4B, respectively, but for replicate 3 of the EdU pulse T chase experiment.

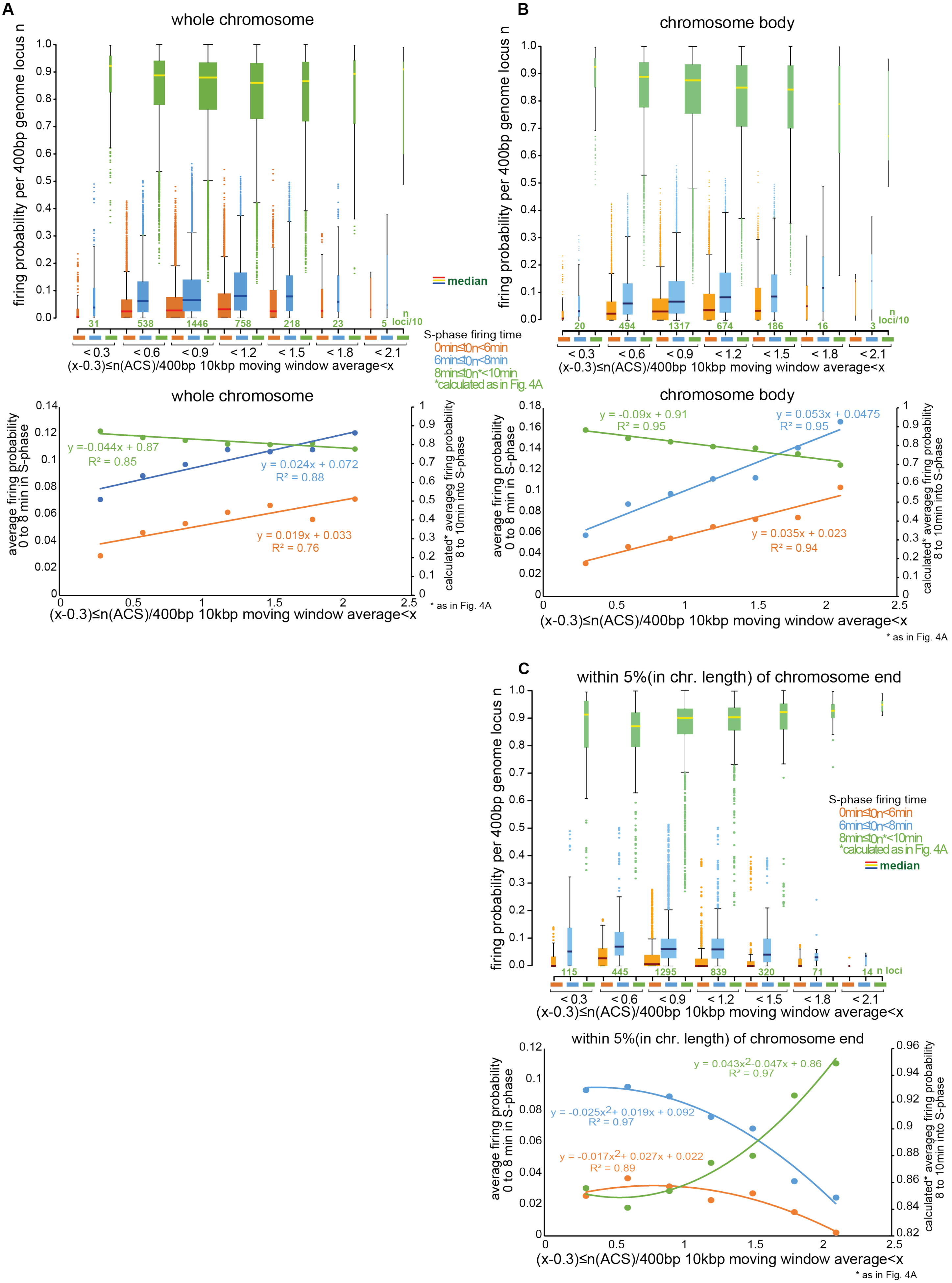


**Fig. S4: Firing probabilities at chromosome ends are inversely correlated to ACS density.**

**A.** The entire yeast genome was divided into bins of ACS density (10kbp moving average of n(ACS)/400bp, see Figs. 5B and 7A), and the boxplot distribution of firing probabilities (defined as in Fig. 7A) in the first half (0 to 6min, orange), and the 3rd (6 to 8min, blue) and last quarters (8 to 10min, green) of S-phase was determined for each bin (**top**). Since origin firing is under detected in the last quarter of S-phase, we used the calculated firing probabilities for the last S-phase quarter (see Fig. 7A). The average firing probability for each bin from each time period in S-phase in the top panel was then plotted against the upper limit of ACS density in each bin, (**bottom**).**B.** As in A. but for 90% of the genome without chromosome ends. **C.** As in A and B but only for chromosome ends. Chromosome ends comprise 10% of the genome and are located within 5% of chromosome length from the left and right end of each chromosome. The end region from the right end of chromosome 3 is extended to 10% of the length of chromosome 3 to include the silent mating type locus HMR.

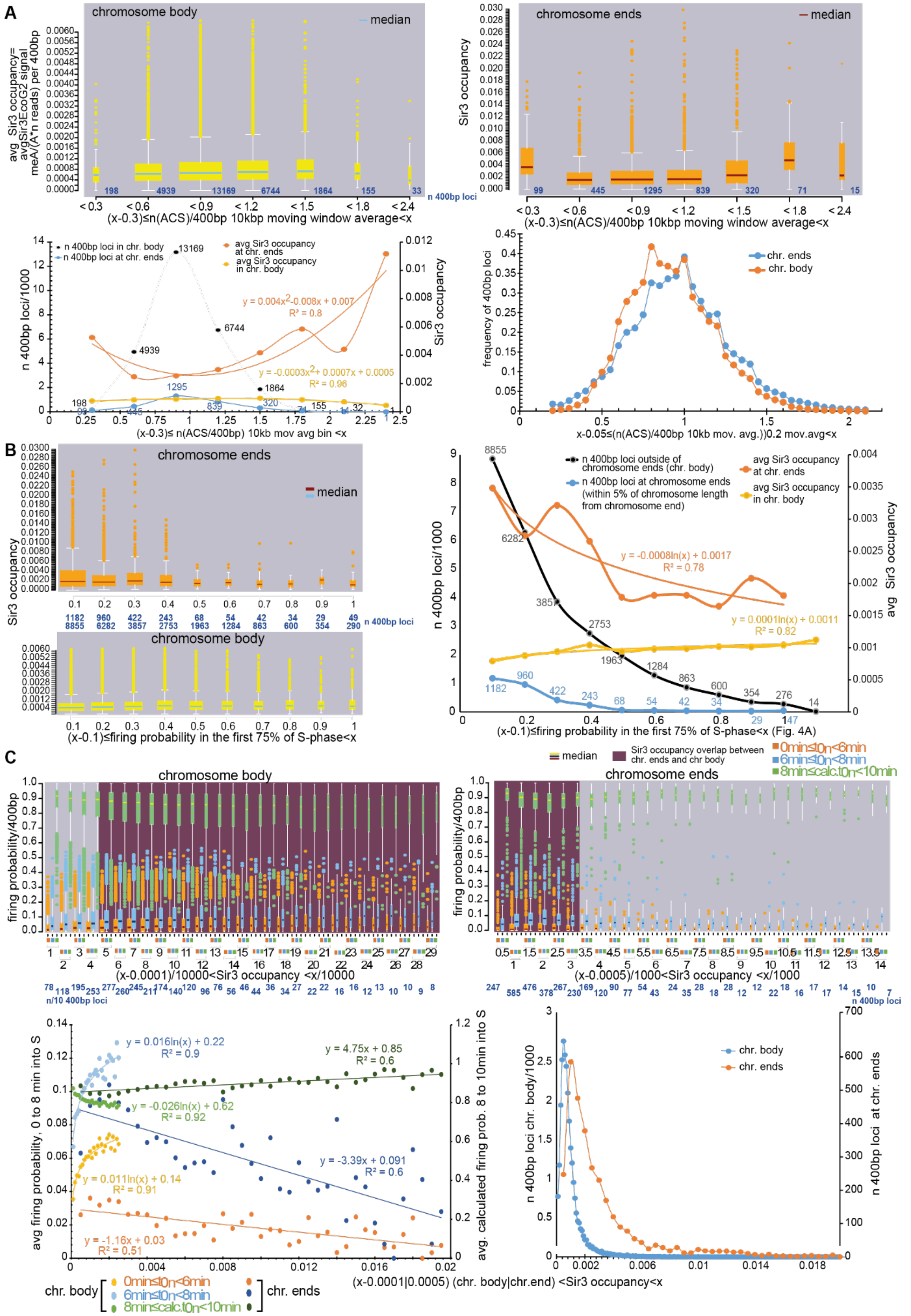
A
chromosome body
median
avg Sir3 occupancy= avgSir3EcoG2 signal meA/(A*n reads) per 400bp
198 4939 13169 6744 1864 155 33 n 400bp loci
< 0.3 < 0.6 < 0.9 < 1.2 < 1.5 < 1.8 < 2.4
(x-0.3)≤n(ACS)/400bp 10kbp moving window average<x
chromosome ends
median
Sir3 occupancy
99 445 1295 839 320 71 15
< 0.3 < 0.6 < 0.9 < 1.2 < 1.5 < 1.8 < 2.4
(x-0.3)≤n(ACS)/400bp 10kbp moving window average<x
n 400bp loci in chr. body
n 400bp loci at chr. ends
avg Sir3 occupancy at chr. ends
avg Sir3 occupancy in chr. body
n 400bp loci/1000
Sir3 occupancy
y = 0.004x2-0.008x + 0.007
R² = 0.8
y = -0.0003x2+ 0.0007x + 0.0005
R² = 0.96
(x-0.3)≤ n(ACS/400bp) 10kb mov avg bin <x
chr. ends
chr. body
frequency of 400bp loci
x-0.05≤(n(ACS)/400bp 10kb mov. avg.))0.2 mov.avg<x
B
chromosome ends
median
Sir3 occupancy
1182 960 422 243 68 54 42 34 29 49
8855 6282 3857 2753 1963 1284 863 600 354 290 n 400bp loci
chromosome body
(x-0.1)≤firing probability in the first 75% of S-phase<x
n 400bp loci outside of chromosome ends (chr. body)
n 400bp loci at chromosome ends (within 5% of chromosome length from chromosome end)
avg Sir3 occupancy at chr. ends
avg Sir3 occupancy in chr. body
n 400bp loci/1000
avg Sir3 occupancy
y = -0.0008ln(x) + 0.0017
R² = 0.78
y = 0.0001ln(x) + 0.0011
R² = 0.82
(x-0.1)≤firing probability in the first 75% of S-phase<x (Fig. 4A)
C
median
Sir3 occupancy overlap between chr. ends and chr body
0min≤t0n<6min
6min≤t0n<8min
8min≤calc.t0n<10min
chromosome body
firing probability/400bp
(x-0.0001)/10000<Sir3 occupancy <x/10000
n/10 400bp loci
chromosome ends
firing probability/400bp
(x-0.0005)/1000<Sir3 occupancy <x/1000
n 400bp loci
avg firing probability, 0 to 8 min into S
avg. calculated firing prob. 8 to 10min into S
y = 0.016ln(x) + 0.22
R² = 0.9
y = 4.75x + 0.85
R² = 0.6
y = -0.026ln(x) + 0.62
R² = 0.92
y = -3.39x + 0.091
R² = 0.6
y = 0.011ln(x) + 0.14
R² = 0.91
y = -1.16x + 0.03
R² = 0.51
chr. body
chr. ends
0min≤t0n<6min
6min≤t0n<8min
8min≤calc.t0n<10min
n 400bp loci chr. body/1000
n 400bp loci at chr. ends
chr. body
chr. ends
(x-0.0001|0.0005) (chr. body|chr.end) <Sir3 occupancy<x

**Fig. S5: The Sir complex delays origin firing at chromosome ends until late S-phase. A.** The entire yeast genome was divided into bins of ACS density (10kbp moving average of n(ACS)/400bp, see Fig. 5B) like in Fig. S4, and bins were separated into a set localizing to the chromosome body (**top left**, defined as in Fig. S4B) or chromosome ends (**top right**, defined as in Fig. S4C). The boxplot distribution of Sir3 occupancy was then determined for each bin. Sir3 occupancy is determined as noted on the y axis of the top left panel, from the *in vivo* Sir3 foot printing experiment with Sir3EcoG2 based on long-read nanopore sequencing (*9*) (Data S3). We used an *in vivo* foot printing dataset instead of the ChIP-seq dataset from(*13*), because *in vivo* foot printing correlates with Sir3 ChIP-seq but is more sensitive and provides better coverage of repetitive sequences on chromosome ends (Data S3). The **bottom left** panel shows the scatterplot between the average Sir3 occupancy per bin of ACS density from the box plot distributions in the top panels and the upper limit of ACS density of each bin. The **bottom right** panel shows the normalized distribution of ACS density in chromosome bodies and chromosome ends. **B.** The genome was divided into bins of origin firing probabilities/400bp in the first 75% of S-phase (the 1$^{st}$ half and the 3$^{rd}$ quarter of S-phase, see 1$^{st}$ and 2$^{nd}$ panels from the bottom in Fig. 7A), and the box plot distribution of Sir3 occupancy (from A) was determined for each bin at chromosome ends (**top left**), or in the chromosome body (**bottom left**). The right panel shows the scatterplot between the average Sir3 occupancy per bin of origin firing probability early in S-phase from the box plot distributions in the left panels, and the upper limit of origin firing probability of each bin. **C.** The genome was divided into bins of Sir3 occupancy and the box plot distribution of measured or calculated origin firing probabilities/400bp in the 1$^{st}$ half, the 3$^{rd}$ (measured) or the last quarter (calculated) of S-phase (see respectively the 1$^{st}$, 2$^{nd}$ and 5$^{th}$ panels from the bottom in Fig. 7A), was determined for each bin in the chromosome body (**top left**) or at chromosome ends (**top right**). The **bottom left** panel shows the scatterplot of the average origin firing probability at different times in S-phase per Sir3 occupancy bin from the box plot distributions in the left panels, and the upper limit of Sir3 occupancy of each bin. The **bottom right** panel shows the distribution of Sir3 occupancy in chromosome bodies and chromosome ends.

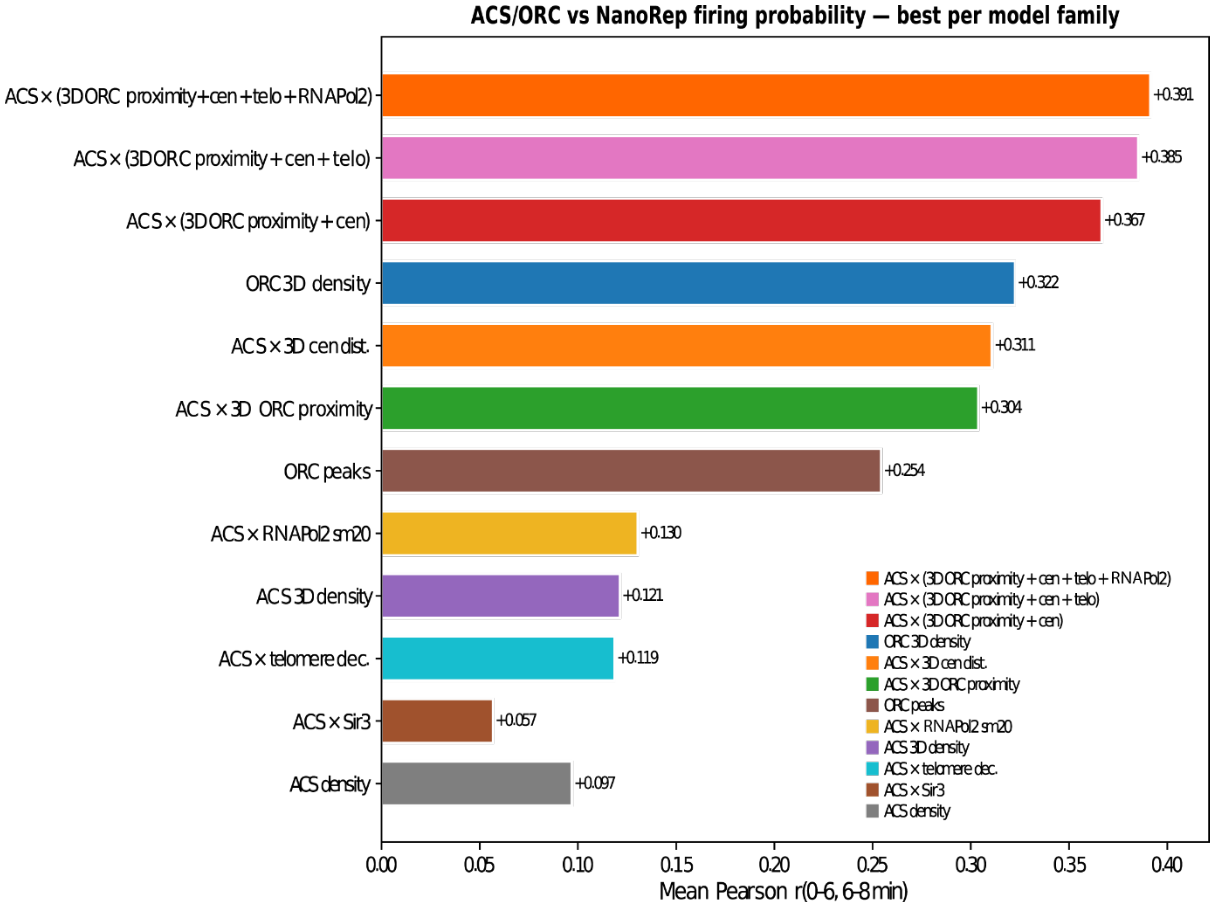


**Figure S6. Ranking of ACS/ORC-based models by correlation with NanoRep firing probability.** The models are divided into two categories. The multiplicative ones (ACS x some feature) *correct* the bare ACS density with one or more factors. The others (e.g. ORC peaks) illustrate the correlation of other profiles with the firing probabilities. Bars are sorted by descending mean Pearson r descending except the ACS density alone, which is considered to be a baseline and it is shown at the bott

From bottom to top :

<u>12. ACS density</u>

The smoothed ACS count per bin (10 kbp Gaussian kernel), no other correction. It constitutes the baseline in this plot.

<u>11. ACS × Sir3</u>

Multiplicative implementation of the Sir3 suppression on the ACS density profile. High Sir3 reduces early firing probability at chromosome ends. Sir3 signal is smoothed at 20 kbp and min-max normalized before multiplication, using the following formula: ACS_density * (1 - 10 * sir3_norm_20kbp). The drop in correlation relative to ACS density alone is probably caused by the mild stimulatory effect of moderate Sir3 levels in chromosome bodies on early origin firing (Fig. S5), which is not factored in the Sir3 suppression function used here.

<u>10. ACS × telomere dec.</u>

Multiplicative implementation of the telomere suppression on the ACS density profile. Bins near chromosome ends are suppressed by an exponential decay of their linear distance to the nearest chromosome end, with the following formula: ACS_density * (1 - 0.87 * exp(-d_telo / 14)), where d_telo = min(bp, chrom_length - bp) in kbp. This is conceptually similar to the Sir3 values, but the suppression function is simplified.

9. ACS 3D density

Replaces each bin's ACS with the contact-weighted average ACS density in its 3D Hi-C neighborhood, from D'Asaro S-phase WT(*7*). The bin's own ACS density is discarded: it inherits the mean ACS of bins that contact it in 3D. The formula used is value for bin i = (sum_j ACS[j] * w_ij) / (sum_j w_ij) where w_ij = ICE-balanced Hi-C contact strength between bin i and bin j.

8. ACS × RNAPol2 sm20

Multiplicative RNAPol2 boost implemented on the ACS density profile. RNAPol2 is the RNAPol2 G1 ChIP-seq signal, smoothed at 20 kbp and normalised. The formula used is ACS_density * (1 + 2 * pol2_norm_20kbp)

7. ORC peaks

Count of ORC ChIP-seq peaks per bin, smoothed at 20 kbp. Same approach as ACS 20 kbp smoothing.

6. ACS × 3D ORC proximity

Multiplicative implementation of the 3D ORC proximity effect on the ACS density. The density is multiplied by (1 + alpha * bin_ORC_proximity), where ORC_proximity = per-chromosome percentile rank of total ICE-balanced contact strength with ORC-bound bins from Hi-C. ORC peaks are pre-smoothed at 20 kbp before computing the ORC flag (same smoothing as ORC peaks bar). The bin's own ACS structure is preserved, just scaled accordingly to the 3d ORC proximity profile.

5. ACS × 3D cen dist.

Multiplicative implementation of the centromere correction via 3D distance. Bins near centromeres in 3D space (from(*10*)) get boosted by an exponential decay, following this formula: ACS_density * (1 + 50 * exp(-d3d / 50)) where d3d = 3D Euclidean distance from bin to centromere (nm)

4. ORC 3D density

Replaces each bin's ORC count with the contact-weighted average of ORCs in its 3D Hi-C neighborhood (from D'Asaro S-phase WT,(*7*)). ORC peaks are pre-smoothed at 20 kbp before the 3D averaging (same smoothing as the ORC peaks bar below). The bin's own ORC is discarded: it inherits the mean ORC of bins that contact it in 3D. The formula used is value for bin i = (sum_j ORC_20kb[j] * w_ij) / (sum_j w_ij) where w_ij = ICE-balanced Hi-C contact strength between bin i and bin j

3. ACS × (3D ORC proximity + cen)

ORC proximity boost and centromere 3D boost applied together additively inside the same factor: ACS_density * (1 + 50 * exp(-d3d/50) + 20 * prox)

2. ACS × (3D ORC proximity + cen + telo)

Adds telomere decay on top of the Full model, with the formula: ACS_density * telo_factor * (1 + 50 * exp(-d3d/50) + 20 * prox), where telo_factor = (1 - 0.87 * exp(-d_telo / 14))

1. ACS × (3D ORC proximity + cen + telo + Pol2)

All four factors combined: centromere 3D boost, ORC Hi-C proximity, telomere decay, and RNAPol2 boost (20 kb smoothed). The best-performing model in the correlation analysis. The final formula is : ACS_density * telo_factor * (1 + 2 * pol2_sm20) * (1 + 50 * exp(-d3d/50) + 20 * prox).

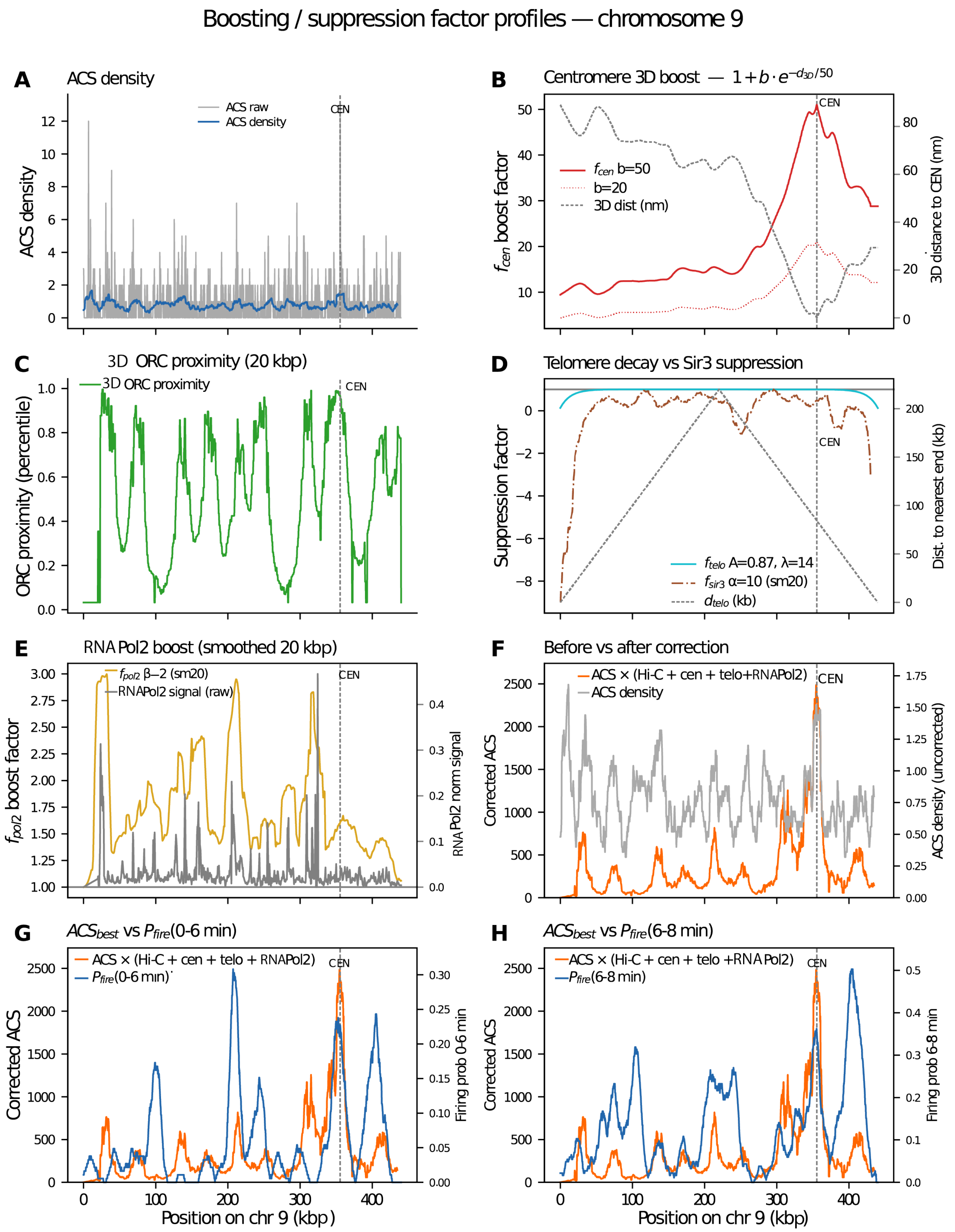


**Figure S7. Illustration of the effect that the changes that the ACS/ORC-based models apply to the ACS density profile on chromosome 9.**
Each panel traces a factor (or combination of factors) used to correct the 10kb smoothed ACS density in Fig. S6, culminating in the best predictive model versus measured firing probability from Fig. 7A.

(A) ACS density at 400 bp resolution: raw counts (light grey) and 10 kb smoothed profile (blue) used as the baseline signal throughout all corrections.
(B) Centromere 3D boost factor: $f\_cen(i) = 1 + b \cdot \exp(-d\_3D(i)/50)$. The 3D distance to the centromere (grey dashed, right axis) modulates a multiplicative boost. Two parameter settings are shown: b=50 (solid red) and b=20 (dotted). Centromere position marked by vertical dashed line.
(C) 3D ORC proximity: the percentile rank of Hi-C contact frequency with the nearest ORC (solid green), used as a proxy for spatial co-localization with origins.
(D) Telomere decay and Sir3 suppression: two independent suppression factors plotted on the same axis. $f\_telo(i) = 1 - 0.87 \cdot \exp(-d\_telo(i)/14)$ (cyan) represents the decay in the negative influence of telomeres on ACS density's effect on early origin firing with increasing linear distance from chromosome ends (grey dashed, right axis). $f\_sir3(i) = 1 - 10 \cdot Sir3\_sm20(i)$ (brown, dash-dot) captures the repressive effect that Sir3 occupancy smoothed at 20 kb has on early origin firing at chromosome ends. Both are bounded at 1.0 (no effect) far from telomeres / Sir3-poor regions
(E) RNAPol2 boost: raw RNAPol2 ChIP signal (grey, right axis) and the resulting smoothed boost factor $f\_pol2(i) = 1 + 2 \cdot Pol2\_sm20(i)$ (gold) applied to the corrected ACS. Only the 20 kb smoothed version ($\beta$=2) is retained as the best-performing parametrization.
(F) Before and after correction: comparison of the uncorrected ACS density (grey) with the fully corrected profile $ACS\_corrected = ACS\_10kb \times f\_cen \times f\_orc \times f\_telo \times f\_RNApol2$ (orange). The corrected profile amplifies ORC-proximal regions in chromosome bodies while suppressing Sir3-rich domains at chromosome ends.
(G-H) Best corrected ACS vs. ground-truth firing probability for 0–6 min ($1^{st}$ half of S-phase, Fig. 7A) (G) and 6–8 min ($3^{rd}$ quarter of S-phase, Fig. 7A) (H). The corrected ACS (orange, left axis) is overlaid on the experimental NanoRep firing probability (blue, right axis). Both axes are scaled independently to fill the panel; the qualitative agreement is reflected by Pearson $r \approx 0.39$ for both time windows.

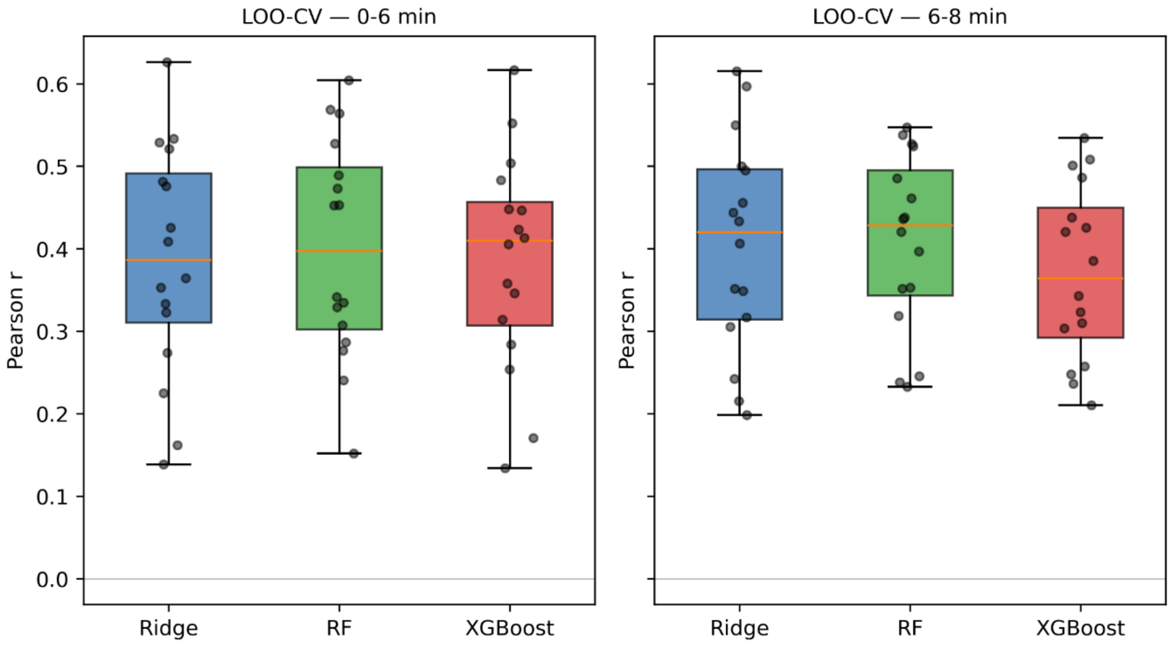


**Figure S8. Leave-one-chromosome-out cross-validation prediction performance for predicting NanoRep firing probability at 4 kb resolution.**
Each panel shows the distribution of per-chromosome Pearson r values (one point per held-out chromosome) for Ridge (α=1.0), Random Forest (max_depth=10, 150 trees), and XGBoost (max_depth=4, 150 trees), for the 0–6min (left) and 6–8 min (right) NanoRep firing probabilities targets (see Fig. 7A and Data S3).
Each model used a total of 12 features at 4 kb bin resolution. They have been trained in a Leave One Chromosome Out cross-validation (LOCO-CV). (*10*)The features are the following (i) ACS density smoothed at 10 kb; (ii) ORC peak count smoothed at 20 kb; (iii) telomere distance; (iv) 3D Euclidean distance to the centromere (*10*); (v–vi) Sir3 occupancy and RNAPol2 ChIP signals, both smoothed at 20 kbp; (vii–viii) ACS × 3D ORC proximity, computed from two Hi-C datasets independently (early S-phase, GSE309730, (*7*); asynchronous wild-type, GSE227687, (*8*)); and (ix–xii) ACS and ORC 3D contact-weighted neighborhood densities (intra-chromosomal only), also from both Hi-C datasets.

In LOCO-CV, Ridge achieves r=0.39 (0–6 min) and r=0.41 (6–8 min), essentially tied with RF (r=0.39 and 0.38) and slightly ahead of XGBoost (r=0.36 for both). The linear model (Ridge) proves as competitive as the non-linear ensemble methods, consistent with a predominantly additive and locally smooth relationship between the 12 features and firing timing.

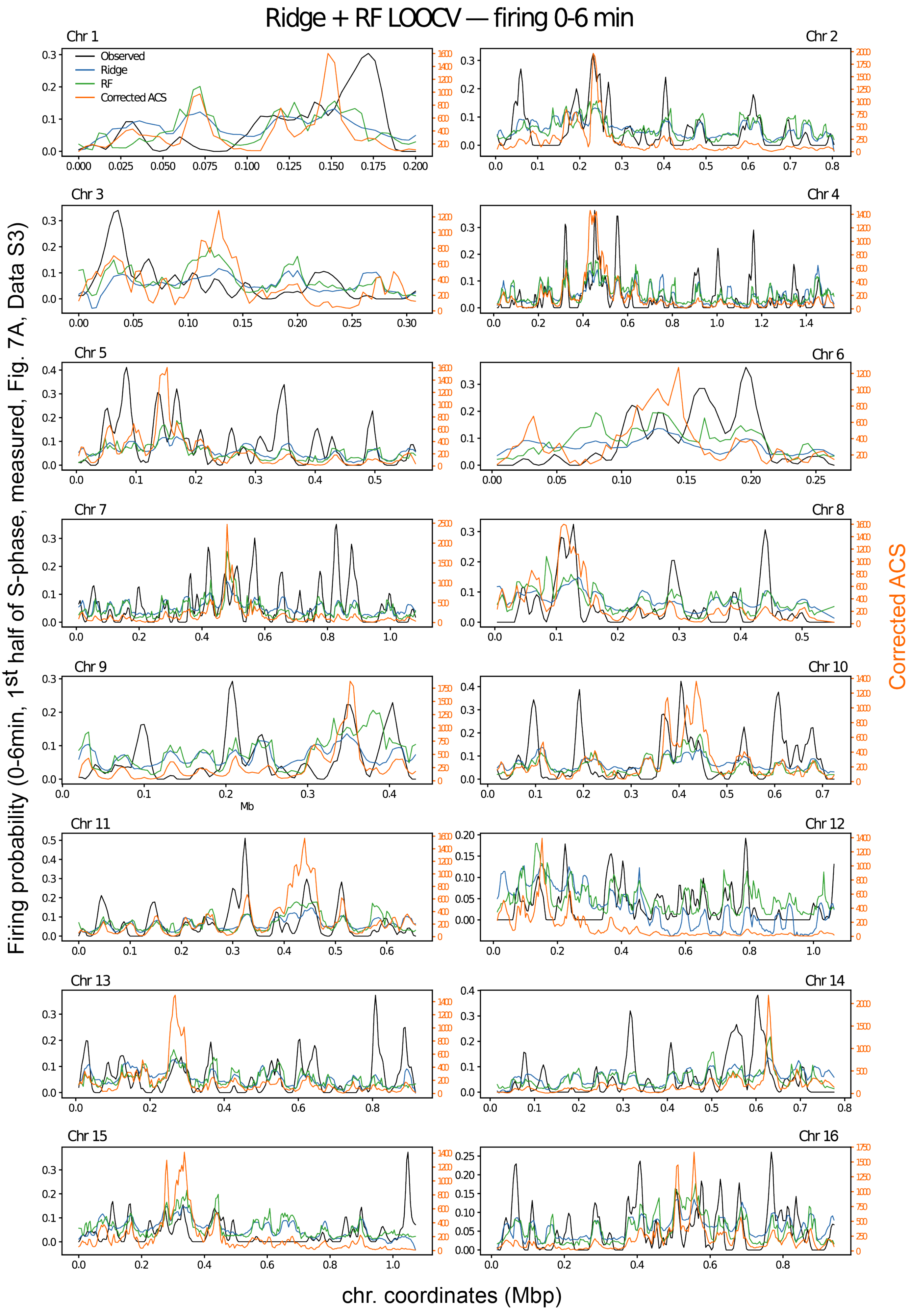
Ridge + RF LOOCV — firing 0-6 min
Chr 1
Observed
Ridge
RF
Corrected ACS
Chr 2
Chr 3
Chr 4
Chr 5
Chr 6
Chr 7
Chr 8
Chr 9
Mb
Chr 10
Chr 11
Chr 12
Chr 13
Chr 14
Chr 15
Chr 16
Firing probability (0-6min, 1st half of S-phase, measured, Fig. 7A, Data S3)
Corrected ACS
chr. coordinates (Mbp)

**Figure S9. Per-chromosome LOOCV profiles of predicted vs observed 0-6min firing probability (in the 1st half of S-phase).** Ridge regression (blue) and Random Forest (green) were trained on 12 features (ACS_10 kb, ORC_20 kb, telomere distance, 3D centromere distance, Sir3_20 kb, RNAPol2_20 kb, ACS × 3D ORC proximity from two Hi-C datasets, and ACS/ORC contact-weighted 3D densities from two Hi-C datasets) using leave-one-chromosome-out cross-validation (LOCO-CV, 16 folds). The best multiplicative ACS correction from Fig. S6 (orange, right axis) is shown for comparison.

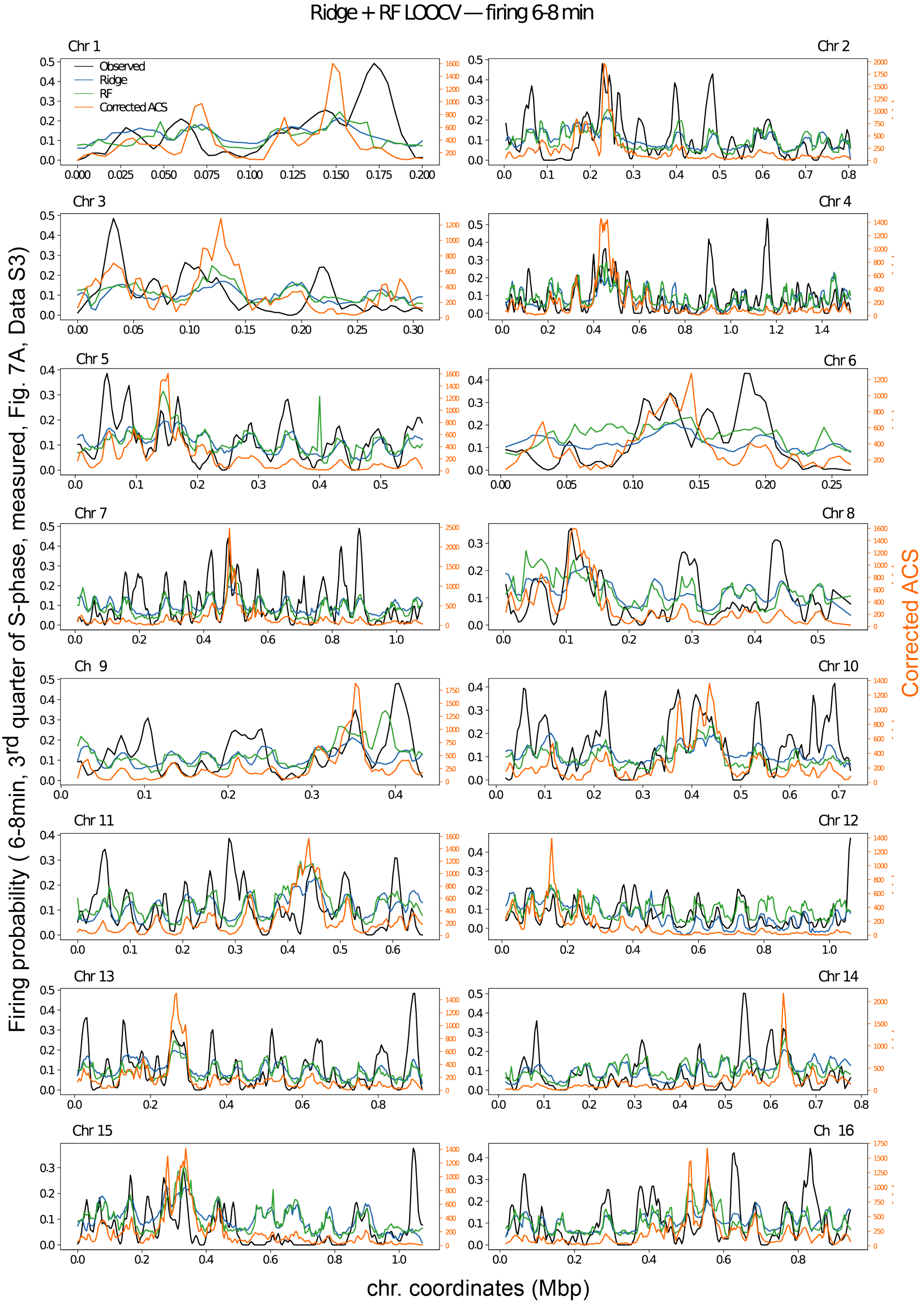
Ridge + RF LOOCV — firing 6-8 min
Observed
Ridge
RF
Corrected ACS
Chr 1
Chr 2
Chr 3
Chr 4
Chr 5
Chr 6
Chr 7
Chr 8
Ch 9
Chr 10
Chr 11
Chr 12
Chr 13
Chr 14
Chr 15
Ch 16
Firing probability ( 6-8min, 3rd quarter of S-phase, measured, Fig. 7A, Data S3)
Corrected ACS
chr. coordinates (Mbp)

**Figure S10. Per-chromosome LOOCV profiles of predicted vs observed 6-8min firing probability.** Ridge regression (blue) and Random Forest (green) were trained on 12 features (ACS_10kbp, ORC_20kb, telomere distance, 3D centromere distance, Sir3_20kbp, RNAPol2_20kbp, ACS × 3D ORC proximity from two Hi-C datasets, and ACS/ORC contact-weighted 3D densities from two Hi-C datasets) using leave-one-chromosome-out cross-validation (LOCO-CV, 16 folds). The best multiplicative ACS correction from Fig. S6 (orange, right axis) is shown for comparison.

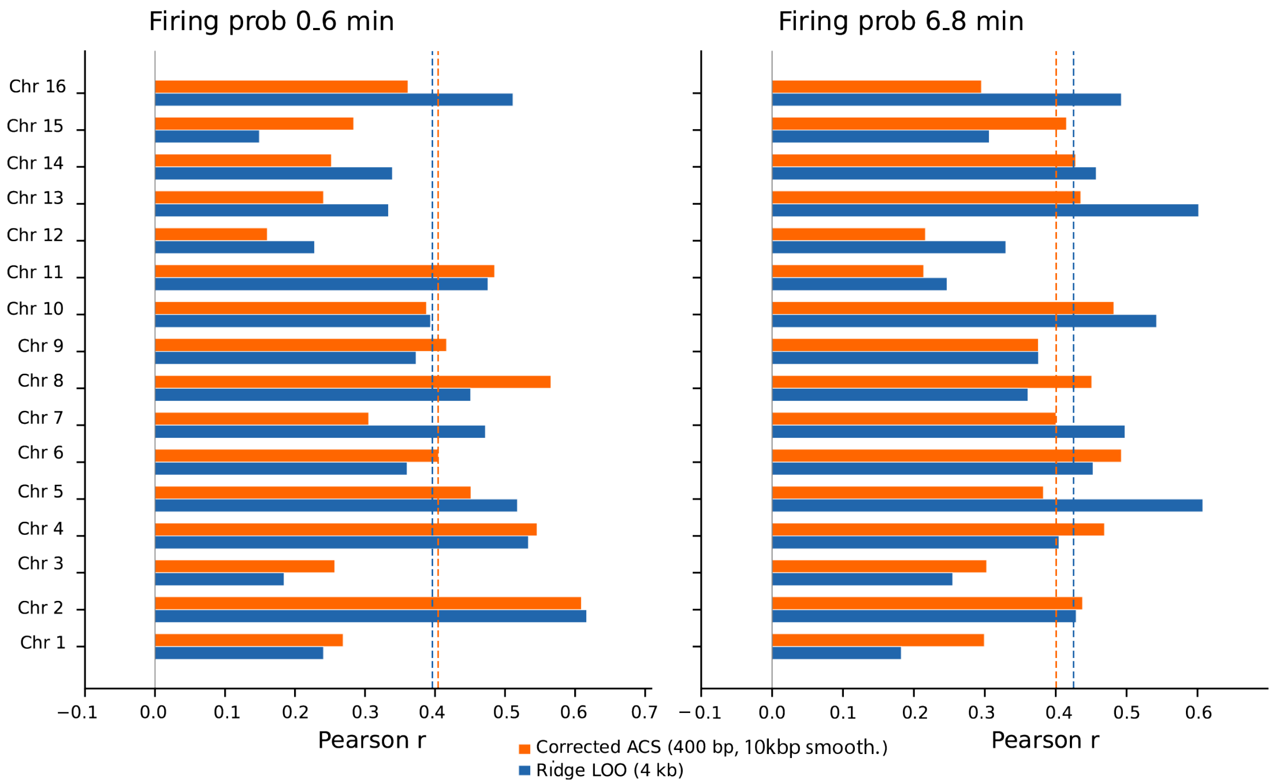


**Figure S11. Per-chromosome Pearson correlation of corrected ACS density and Ridge LOO predictions with NanoRep firing probability.** Left: 0–6 min (1st half of S-phase) firing window; right: 6–8 min (3rd quarter of S-phase). Orange bars: corrected ACS (Full + telomere decay + RNAPol2 sm20 β=2) at 400 bp native resolution. Blue bars: Ridge regression LOO predictions (12 features, Z-scored per fold) at 4 kb resolution. Chromosomes are ordered naturally (1–16, top to bottom). Dashed vertical lines mark the global (all-chromosome combined) Pearson r for each method (orange: corrected ACS; blue: Ridge). Despite the different resolutions (400 bp vs 4 kb) and model classes (hand-crafted multiplicative vs machine learning), per-chromosome r values are broadly consistent between the two approaches, with most chromosomes clustering between r = 0.2 and r = 0.6 and a few showing substantial divergence (e.g., chr16 0–6 min: ACS r = 0.36, Ridge r = 0.51). The near-identical global r (~0.40 for both targets) confirms that the simple multiplicative correction captures the same linearly predictable fraction of firing-probability variance as the full 11-feature Ridge model.

**Data S1. (separate file)**

EdU/T peaks with corresponding fork velocities (fv) and firing time ($t_{0n}$) from all rep1 and rep 2 EdU pulse and T chase datasets (see Fig. S2). This file contains the tabulated data behind the figures 1, 2, 3D-I and 4A.

**Data S2. (separate file)**

All 400bp origin loci from all rep1 and rep 2 EdU pulse and T chase datasets (see Fig. S2). This file contains the tabulated data behind the figures 3A-C.

**Data S3. (separate file)**

Probability of Origin Firing during S-phase determined from the genome wide distribution of firing time $t_{0n}$ per 400bp bin, described in Fig. 4A.

**Code file (separate file)**

**NanoRep_pipeline.tar** contains Perl scripts for EdU/T peak identification on single reads and calculations of firing times and fork velocities per peak.
